# Does the visual word form area split in bilingual readers? A millimeter-scale 7T fMRI study

**DOI:** 10.1101/2022.11.10.515773

**Authors:** Minye Zhan, Christophe Pallier, Stanislas Dehaene, Laurent Cohen

## Abstract

In expert readers, a brain region known as the visual word form area (VWFA) is highly sensitive to written words, exhibiting a posterior-to-anterior gradient of increasing sensitivity to orthographic stimuli whose statistics match those of real words. Using high-resolution 7T fMRI, we ask whether, in bilingual readers, distinct cortical patches specialize for different languages. In 21 English-French bilinguals, unsmoothed 1.2 mm fMRI revealed that the VWFA is actually composed of several small cortical patches highly selective for reading, with a posterior-to-anterior word similarity gradient, but with near-complete overlap between the two languages. In 10 English-Chinese bilinguals, however, while most word-specific patches exhibited similar reading specificity and word-similarity gradients for reading in Chinese and English, additional patches responded specifically to Chinese writing and, surprisingly, to faces. Our results show that the acquisition of multiple writing systems can indeed tune the visual cortex differently in bilinguals, sometimes leading to the emergence of cortical patches specialized for a single language.

## Introduction

Half of humanity speaks more than one language, and many adults can read more than one language and master multiple writing systems. How does the visual cortex accommodate the recognition of written words in two languages, possibly using two different scripts? Much previous research has shed light on the mechanisms of reading acquisition in a single script. Within the left ventral occipito-temporal cortex (VOTC), the recognition of written words mobilizes a small cortical area which has been termed the visual word form area (VWFA) (Cohen et al., 2000). This region emerges during reading acquisition (Brem et al., 2010; Dehaene et al., 2010, 2015; Dehaene-Lambertz et al., 2018; Feng et al., 2020, 2022; Schlaggar & McCandliss, 2007) and becomes attuned only to the script that the person has learned to read (Baker et al., 2007; Cohen & Dehaene, 2004; Dehaene et al., 2015; Dehaene & Cohen, 2011; Szwed et al., 2014). In readers of all languages, the VWFA occupies a reproducible location within a mosaic of cortical regions specialized for the recognition of various categories of visual stimuli such as faces, bodies, objects or places. This reproducible specialization is thought to be based on a combination of factors including foveal bias (Hasson et al., 2002), preference for geometrical features (Long et al., 2018; Srihasam et al., 2014; Szwed et al., 2009) and preexisting connectivity to distant language areas (Bouhali et al., 2014; Feng et al., 2022; Hannagan et al., 2015; Mahon & Caramazza, 2011; Saygin et al., 2016). Longitudinal studies of school children show that the VWFA acquires its specialization for written words within the first few months of schooling (Dehaene-Lambertz et al., 2018). Its lesion, in literate individuals, results in pure alexia, a selective reading impairment (Cohen et al., 2016; Gaillard et al., 2006; Pflugshaupt et al., 2009).

How populations of neurons in the VWFA encode written words is not known (for proposals, see Hannagan et al., 2021; Rajalingham et al., 2020), but one of its key macroscopic features is a sensitivity to the statistics of reading: the response of the VWFA increases as letter strings increasingly respect the statistical distribution of letters in real words, with an increasing gradient of sensitivity along the posterior-to-anterior axis along the VOTC (Binder et al., 2006; Dehaene et al., 2005; Vinckier et al., 2007; Woolnough et al., 2021). Thus, it is plausible that, during reading acquisition, neurons in the VWFA internalize the statistics of letters and their combinations. Due to the limited spatial resolution of imaging methods, however, which frequently smooth and average data across many individuals, it is not known whether the macroscopic gradient along the VOTC results from a continuous increase in sensitivity to overall word similarity, or from a chain of discrete patches of cortex, each possibly responsive to a hierarchically higher-level orthographic component such as letters, bigrams, and larger chunks of letters (Dehaene et al., 2005), in part through interactive bottom-up and top-down influences (Woolnough et al., 2021).

Here, we ask how plasticity allows this architecture to adapt to reading in two different languages in bilingual readers. Do distinct cortical patches implement reading in different languages? The above statistical learning hypothesis leads to contrasted predictions depending on whether the two languages use the same alphabet (e.g. English and French), or two widely dissimilar scripts, typically alphabetic vs. logographic (e.g. Chinese). Whenever the two languages use the same alphabet, words share similar visual features in both languages, and the visual cortex projects to the same distant language areas (Perani et al., 1998; Xu et al., 2017). One may therefore predict that the same cortical patches would be used in both languages. Local patches of visual cortex would compile letter statistics without any consideration of which language they transcribe, and should therefore be sensitive to the overall orthographic statistics, pooled across both languages (in proportions that may vary depending on the dominance of one language over the other for a given individual). Indeed there is currently no imaging evidence of language-specific patches in the VOTC, either between monolingual (Rueckl et al., 2015) or within bilingual individuals (Brignoni-Perez et al., 2020; Hsu et al., 2015; Jamal et al., 2012; Li et al., 2021; Meschyan & Hernandez, 2006). Accordingly, we know of no reports of developmental dyslexia or pure alexia differently affecting two languages using the same alphabet (Wechsler, 1977). Nevertheless, it remains possible that distinct cortical patches or columns specialize for one or the other language, and incorporate the orthographic statistics of only one language. Such fine-grained specialization may have escaped the relatively coarse spatial resolution of previous 3T fMRI studies, especially using group-level averaging, and the still coarser grain of brain lesions.

Conversely, in bilingual readers of an alphabetic and a logographic script, written words differ in number of learned characters, visual complexity, component shapes and overall contour area. Those factors make it more likely that the VOTC should develops script-specific cortical patches (Srihasam et al., 2014), with each patch encoding the orthographic regularities of a single language. Still, statistical analyses suggest that all scripts rely on similar statistics of line junctions (Changizi et al., 2006; Changizi & Shimojo, 2005). It may therefore still be useful for the visual system to share the same cortical resources between two distinct reading systems. The existing brain imaging data, however limited, mostly support this second possibility. Neither between monolinguals (Rueckl et al., 2015) nor within bilinguals (Xu et al., 2017) is there any strong evidence of distinct script-selective regions. However, multivariate techniques can successfully decode Chinese from English words in the left VOTC (Xu et al., 2017), which may point to specialization beyond the usual resolution of fMRI. In Japanese readers, who master both the syllabic Kana and the logographic Kanji scripts, a double dissociation has been reported between two types of pure alexic patients, with a predominant deficit either for Kana or for Kanji (Paradis et al., 1985; Sakurai et al., 2006, 2008; Sugishita et al., 1992). In developmental dyslexia, a single dissociation between English (impaired) and Japanese (preserved) is on record (Wydell & Butterworth, 1999). Thus, the existence of script-specific cortical patches is a plausible hypothesis, whose empirical assessment requires high-resolution individual brain imaging.

Here, we study the organization of visual word recognition in the VOTC of bilingual participants, while maintaining high spatial precision by using individual analyses of high-resolution 7 Tesla fMRI (1.2 mm isotropic voxels). We first imaged 21 bilingual readers of English and French, two languages written with the same alphabet but different orthographic statistics, then 10 bilingual readers of English and Chinese, two languages which rely on alphabetic and logographic scripts with very different visual features. In each case, following up on our previous work (Vinckier et al., 2007), we created a hierarchy of stimuli with increasing similarity to real words. This design allowed us to assess the cortical implementation of orthographic statistics in three different languages, to look for cortical patches specialized for orthographic components of alphabetic versus logographic scripts, to assess the existence of specialization for one of two available languages, whether they share the same script or not, and to study the modulation of those findings by language dominance.

## Results

### Experiment 1: Reading in English-French bilinguals

#### Localizer for visual categories and overall reading circuit

We recruited 21 English-French bilingual readers (7 dominant in English, 7 fully bilingual, and 7 dominant in French). Participants were first imaged using a localizer for various visual categories, including words in both languages. They passively viewed blocks of English or French words, Arabic numbers, false font strings, faces, bodies, houses, tools and checkerboards, while they were asked to detect an occasional star in order to keep their attention focused (**Figure 1A**).

**Figure 1.**
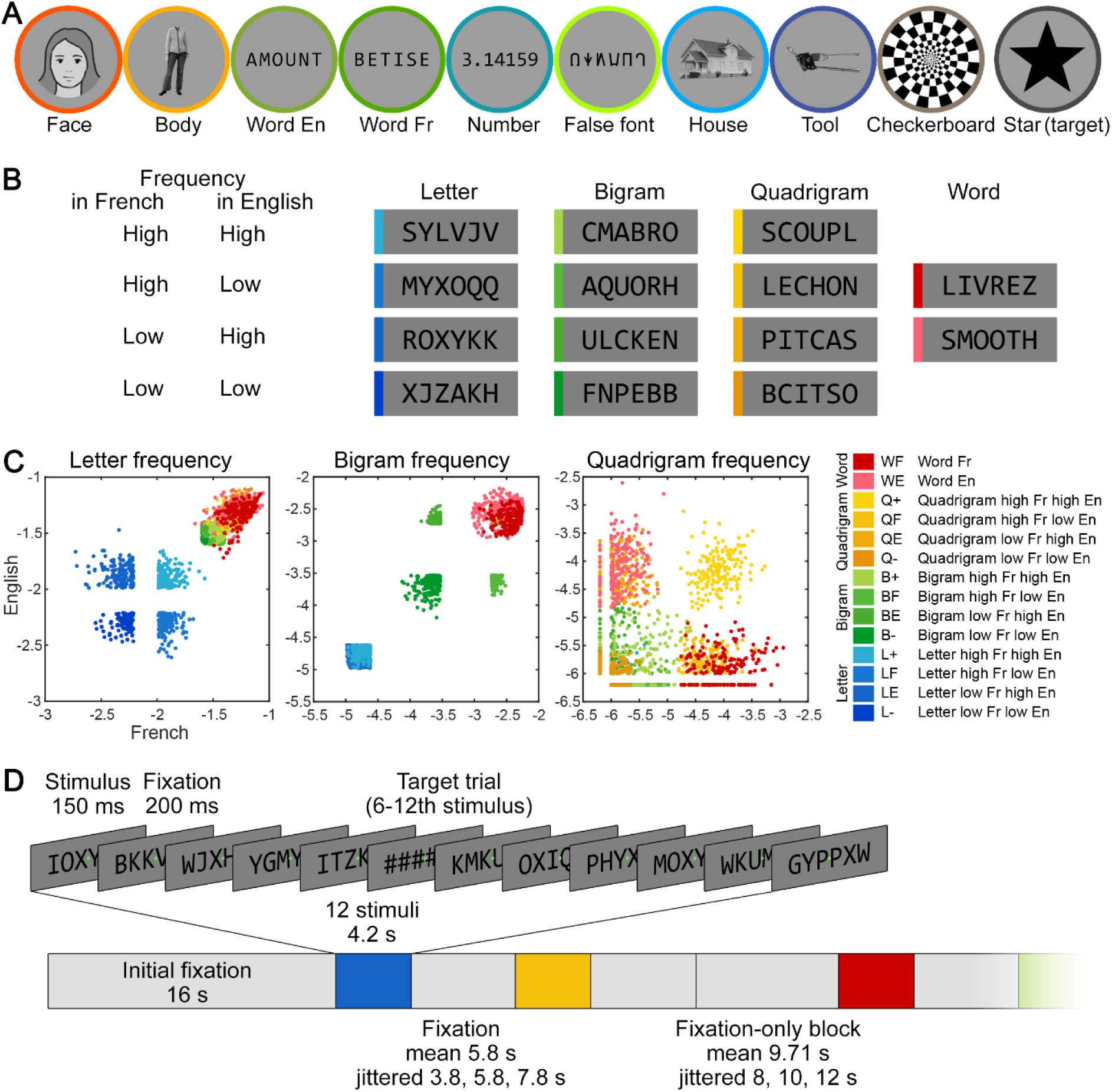
Stimuli and procedure for experiment 1 with English-French bilingual readers. **A.** Examples of the 9 categories of visual stimuli, plus the target, used in the localizer. The actual face stimulus is replaced by an illustration here. **B.** Examples of the 14 categories of visual stimuli in the main fMRI runs, consisting of 6-letter strings whose similarity to words was systematically manipulated. The frequency of letters, bigrams, and quadrigrams, plus the effect of lexicality, were manipulated orthogonally in English and French, resulting in 12 types of non-word letter strings, plus English and French words. **C.** The frequency of letters, bigrams, and quadrigrams in English (y axis) and in French (x axis) for all 14 categories of stimuli (2520 stimuli in total, one dot indicates one stimulus; unit = log10 count per million). The English and French real words (WE and WF) were matched with the corresponding high-frequency quadrigram stimuli (QE and QF) in all 3 parameters. **D.** Organization of an fMRI run, consisting in an alternation of fixation periods and homogeneous blocks of fast stimuli presentation.

We first examined whole-brain word-specific activation at the group level. To facilitate inter-subject averaging, data were smoothed using a Gaussian filter with full-width at half-maximum of 6 mm. We examined the contrast of English & French words > faces, bodies, houses, tools, with a cluster size threshold corrected for multiple comparisons using Monte-Carlo simulations (p<0.001, alpha<0.05, resulting cluster size>74 functional voxels, 1.2 mm isotropic). We found several word-specific clusters along the bilateral superior temporal sulci (STS) and inferior frontal gyri (IFG), with left predominance. In the left VOTC, we did not find the usual VWFA (around Talaraich [TAL] coordinate Y=-56), but found a more anterior cluster in the left occipito-temporal sulcus (OTS), peaking at Y=-29 (**Figure 2A**). Group-level comparisons of English and French words only revealed two small clusters in the right cuneus and vmPFC (alpha<0.05, cluster size>27 functional voxels) that were activated more by English than by French words. The opposite contrast showed no significant activation.

**Figure 2.**
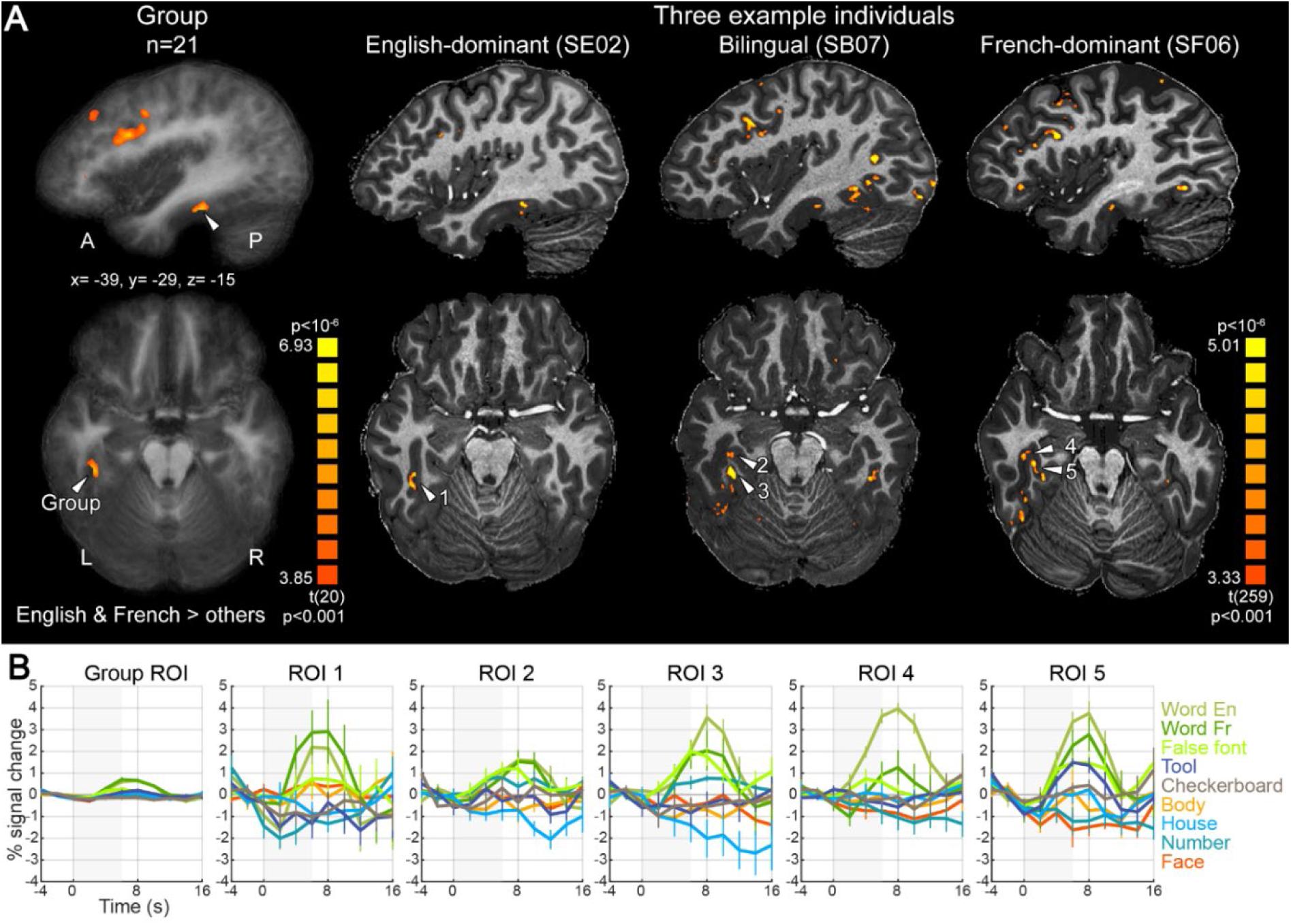
Activation to written words in individual English-French bilingual readers, illustrating the need for single-subject analyses. **A.** In ventral occipito-temporal cortex (VOTC), the group-level contrast of words minus faces, bodies, houses, and tools showed only one anterior word-specific cluster (TAL Y=-29, column 1). However, each individual participant actually showed robust word-specific cortical clusters/patches around the usual location of the VWFA, as illustrated in one participant per language group (columns 2-4; See **Figure S2** for activation in all 21 participants). Overlap between participants was reduced due to individual variability and the small size of those clusters/patches. **B.** Averaged time courses for each category of stimuli in the clusters marked by white arrows in **A**. The shaded area represents stimulus block duration. Error bars denote SEM. Note that the seemingly higher activity in ROI 4 for English than French words (p=0.0063) did not survive whole-brain cluster thresholding (p<0.001), and was not replicated in the main fMRI runs (p=0.434).

As described in a previous study (Glezer & Riesenhuber, 2013), the absence of a VWFA cluster in the group analysis likely resulted from individual anatomical and functional variability. As compared to 3T data, 7T images have enhanced white-gray matter contrast, and the activated clusters/patches are smaller (diameters usually were within 10 mm), largely confined to the gray matter, and with little overlap across participants. Thus, classical brain-wide group analyses are largely inoperative. In contrast, in every single participant, without data smoothing, we observed focal word-specific clusters along the gray matter of the inferior occipital sulcus (IOS) and OTS in the vicinity of the VWFA, showing robust activation time courses (p<0.001 uncorrected, **Figure 2BC**, **Figure S2**). The size of those clusters changed only minimally when using a more stringent voxelwise threshold of p<0.0005. Therefore, all of our subsequent analyses were based on the signals extracted from 773 single-subject word-specific cortical patches, followed by group-level statistical inferences.

In 17 out of 21 participants, word-specific clusters in the VOTC were bilateral, and the remaining 4 participants only showed left-hemisphere clusters. The mean number of activated clusters and voxels was much larger in the left than the right hemisphere (paired t tests, mean of 10.8 vs 3.5 clusters per participant, p=7.50×10^-10^; mean of 853 vs 192 voxels, p=4.58×10^-7^). Around the fusiform gyrus, clusters were mostly dispersed along the IOS and OTS (**Figure S2**), and did not follow any obvious grouping pattern. Again, there were more clusters around the left than the right fusiform region (paired t test, mean of 6.4vs 1.1 clusters per participant, p=4.88×10^-10^; mean of 433 vs 42 voxels, p=9.88×10^-7^).

Previous research has emphasized the similarities between word and face recognition (e.g. Dehaene et al., 2015; Feng et al., 2022; Hasson et al., 2002; Puce et al., 1996). Indeed, faces also require foveal processing and elicit category-specific activation in the VOTC, mesial and adjacent to word-specific activations but with right-hemispheric predominance (Kanwisher et al., 1997; Kanwisher & Yovel, 2006). Parallel to what we did for words, we identified face-specific clusters in the VOTC (contrast: faces > English & French words, bodies, houses, tools, p<0.001 uncorrected, **Figure S2**). We found bilateral VOTC clusters in all participants, in the right more than in the left hemisphere (paired t tests, 6 vs 4.05 clusters, p=0.03; 368 vs 171 voxels, p=4.14×10^-5^). Their location roughly followed a posterior to anterior clustering pattern reproducible across participants, corresponding to the previously described occipital face area (OFA), fusiform face area (FFA), and anterior face patch (AFP) (Tsao et al., 2008). Some face-specific clusters were adjacent to word-specific clusters, e.g. on opposite banks of the same sulcus (see e.g. SB03 and SF05 in **Figure S2**), and occasionally they overlapped by a few voxels.

For both word- and face-specific clusters, we observed anterior activation in the fusiform and OTS regions (Talairach Y coordinate -50 to -22). These anterior clusters are often lacking in fMRI studies of the VWFA, but were reported in intracranial studies (e.g. Lochy et al., 2016; Nobre et al., 1994). We consistently found them here because early in the experiment we optimized the placement of the acquisition slices and improved the signal-to-noise ratio (SNR) affected by the signal dropout around the ear canals (Wandell, 2011). This signal dropout was perceivable in the very first 3 participants (**Figure S1**), although the artifact was anterior and did not affect the VWFA. The SNR there was considerably improved in all subsequent participants.

We also used the localizer to search for significant activation differences between English and French words. We did not find any consistent language-specific activation (within-subject threshold p<0.001 uncorrected). Only 6 participants show some putative language-specific clusters, all outside the VOTC, spread in left and right supramarginal gyri, STS, intraparietal sulcus (IPS), and lateral frontal areas (inferior frontal sulci/gyri: IFS/IFG). However, the language preference of these clusters was not reproduced in the main fMRI runs described below, and was therefore not analyzed further.

Due to the lack of language-specific clusters, we used the bilingual word-specific contrast (English & French words > faces, bodies, houses, tools) to define ROIs for subsequent analyses. To further exclude the possibility that we may have missed language-specific clusters by this bilingual word-specific contrast, we further examined the voxels of two additional language-specific contrasts: English words > faces, bodies, houses, tools; and French words > faces, bodies, houses, tools; both p<0.001 uncorrected, cluster size threshold >4 voxels). The voxels in the language-specific contrasts were mostly overlapping with the ones in the bilingual word-specific contrast (English: mean=91.9% overlapping with bilingual voxels, SD=11.3%; French: mean=86.1%, SD=17.0%, and showed even higher overlap within VOTC (English: mean=958%, SD=5.2%; French: mean=94.8%, SD=7.3%).

Supplementary analysis of the main fMRI runs again failed to show reproducible language specificity: even when voxels were isolated based on their apparent responsivity to a single language in the localizer (e.g. English but not French), the language specificity for the two languages was not consistent across conditions, and was not reproducible between the localizer and the main fMRI runs (see Supplementary Text and **Figure S3**).

#### A hierarchy of stimuli separately manipulating the statistics of English and French

In the main fMRI runs, participants passively viewed mini-blocks of twelve consecutive 6-letter strings presented at a fast rate (stimulus onset asynchrony = 350 ms, see Materials and Methods). Their task was to detect an occasional string of hashtags (######) that occurred randomly in 40% of the blocks. We adopted a design similar to Vinckier et al. (2007), but adapted to bilinguals, by carefully generating pseudowords that orthogonally varied the frequency of a given orthographic component in English and in French. For the letter level, for instance, we generated stimuli whose average letter frequency was low in both languages; or low in one, but high in the other; or high in both – thus resulting a 2×2 design with low/high frequency in English × low/high frequency in French. By manipulating letters, bigrams and quadrigrams in this manner, we obtained 12 categories of pseudoword stimuli increasingly similar to real words, crossing English/French × high/low frequency × letters/bigrams/quadrigrams, to which we further added 2 sets of real English and French words (see **Figure 1B-D**, Materials and Methods). Each letter string was presented only once in the experiment.

We defined 773 bilingual word-specific ROIs in individual participants using the localizer data (p<0.001 uncorrected, cluster size > 4). We checked that almost all of these ROIs also showed above-baseline activity for words (English + French) in the main fMRI runs (749/773 ROIs, including 294/299 VOTC ROIs). We then extracted the activity values (beta, equivalent to % signal change) for the 14 conditions of the main fMRI runs and averaged them across voxels in each ROI.

#### A spatial gradient of word similarity

We first assessed the overall effect of similarity of the stimuli to real words, pooling across both languages without assuming language specificity, by assigning weights to the 14 conditions, ranging from 1 for strings with low-frequency letters in both French and English to 10 for real words. We then normalized these weights between 0 and 1, and used them to fitted a linear regression to the activity per condition within each ROI (See Materials and Methods, and **Figure 3**). Whenever we use the term “slope”, we will be referring specifically to this regression coefficient quantifying the word similarity effect. A significant slope in an ROI would reveal a word similarity effect in that ROI.

**Figure 3.**
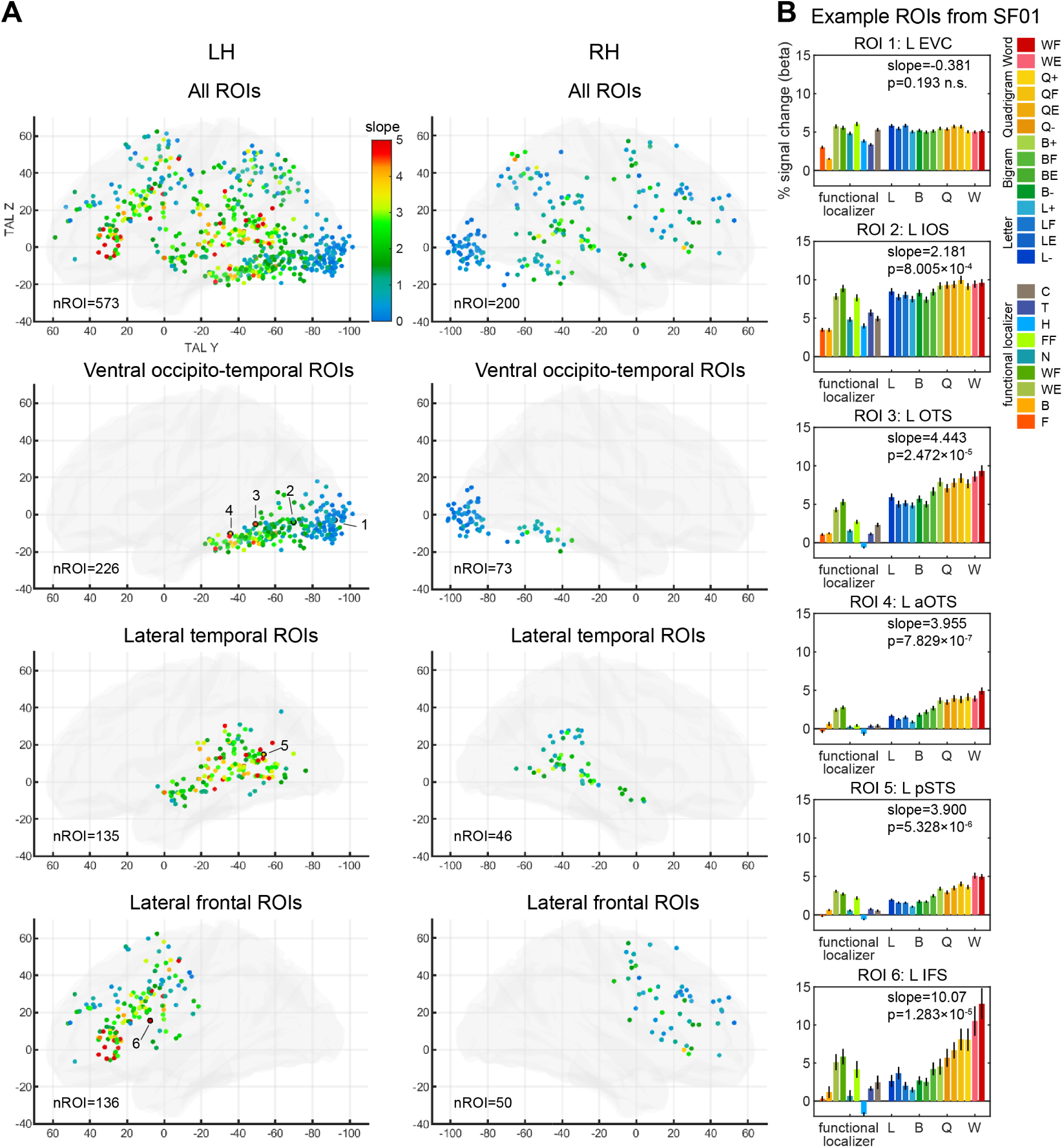
The word similarity effect in English-French bilingual readers. **A.** Word similarity was modeled as the slope of a linear regression over the 14 types of stimuli ranging from low-frequency letters to real words (see Materials and Methods for details). Transparent hemispheres (LH, RH) show the slope value (color-coded) in each of the 773 word-specific ROIs defined from the localizer. There is a posterior-to-anterior gradient of increasing word similarity effect in the VOTC, and two other gradients in the STS and the lateral frontal cortex. **B.** Activity profile in 6 ROIs of a representative participant (SF01; see panel A for ROI localization). Bars represent the activity of each condition in the localizer (left) and the main fMRI runs (right; see color legend at right; note that color codes in **A** and **B** are unrelated). Error bars denote SEM across voxels within each ROI. Abbreviations: EVC: early visual cortex; IOS: inferior occipital sulcus; aOTS: anterior occipito-temporal sulcus; pSTS: posterior superior temporal sulcus; IFS: inferior frontal sulcus.

Among the 773 word-specific ROIs identified across all participants, 64.5% (499/773) showed a significant slope (p<0.05, FDR q<0.05, **Figure 3A top row**). ROIs were grouped into 3 broad anatomical regions (VOTC, lateral temporal, lateral frontal, **Figure 3A rows 2-4**). Inspecting bilateral VOTC, we observed a posterior-to-anterior increase in the slope of the word-similarity effect along the VOTC (**Figure 3**, second row), similar to the spatial gradient observed in (Vinckier et al., 2007). We use the term “gradient” here to denote the gradual changes of a functional property across cortical space. In left VOTC, the gradient of word similarity was highly reproducible across participants: the slope of the word similarity effect was positively correlated with the TAL Y coordinate of the clusters in 19/21 participants (all individual p < 0.028; group-level t test of the regression coefficient against 0, mean coefficient=0.0443, t(20)=16.190, p=5.84×10^-13^). When restricting the analysis to ROIs located around the left fusiform gyrus, thus excluding the early visual cortex ROIs, the gradient was still significant (VOTC ROIs anterior to the posterior collateral sulcus, mean coefficient=0.0444, t(20)=7.101, p=6.98×10^-7^).

In right VOTC, the gradient was significant too (mean coefficient=0.0252, t(13)=3.301, p=0.0057), but coefficients were smaller than in the left hemisphere (paired t test for participants with at least 3 right-hemisphere ROIs, t(13)=2.602, p=0.022), thus showing a left-predominance. While we focus here on VOTC, we also note that there was a trend for similar bilateral gradients in the lateral temporal ROIs along the STS, and in the lateral frontal ROIs around the IFG, although their gradient orientations were different (anterior to posterior, and superior-posterior to inferior-anterior, respectively).

The left VOTC gradient could also be confirmed by comparing words versus control stimuli in the localizer. Activity evoked by false fonts and numbers decreased from posterior to anterior VOTC ROIs (negative regression coefficient values when fitting TAL Y coordinates to the activity, **Figure S4AB**), but this decrease was much less pronounced for English and French words (mean regression coefficients: false fonts=-0.0425, numbers=-0.0484, French words=-0.0082, English words=-0.0111; paired t-tests: all p < 2.4397×10^-10^). The difference between words and control stimuli became massive in anterior ROIs (TAL Y > -50).

We also computed a selectivity index for words versus other categories, separately for left and right ventral ROIs, as [word activity - other activity]/[word activity + other activity], where word activity was the average activity evoked by English and French words, while other activity was the average activity to faces, bodies, houses, and tools. The activity of all conditions was padded with the value of the condition with the most negative beta value, so that all resulting beta values were positive and the resulting selectivity index would range between -1 and 1 (Simmons et al., 2007). In left ventral ROIs, the word selectivity index increased monotonically with the TAL Y coordinate (**Figure S4C**, mean regression coefficient=0.005, one-sample t-test against zero, p=9.52×10^-10^), and continued to increase towards 1 anterior to the VWFA (TAL Y around -56).

In summary, two independent analyses revealed a clear cortical gradient of increasing word selectivity from posterior to anterior VOTC.

#### Dissecting the word similarity effect

We next examined in detail the properties of the word similarity effect in the VOTC (Vinckier et al., 2007). First, we asked whether the word-similarity effect is an exclusive property of word-specific clusters, or whether it reflects a general feature of the VOTC as a whole. We computed the word similarity slope separately within word-specific and face-specific clusters (the latter while excluding voxels overlapping with the word-specific clusters). We found that while 51% of bilateral VOTC word-specific clusters showed a significant slope of the word similarity effect (153/299 ROIs at FDR q<0.05), this was the case for only 9.8% of face-specific clusters (26/264 ROIs at FDR q<0.05). Most of the latter ROIs were in the left hemisphere (23/26 ROIs) and were located around the OTS, IOS, posterior collateral sulcus and mid fusiform gyrus, and 12 of them had 1 to 14 overlapping voxels with word-specific clusters. This analysis indicates that sensitivity to word similarity exists almost exclusively within discrete word-specific cortical patches and their close vicinity, and is not a widespread feature of the entire VOTC: the broad, quasi-continuous extension of the word similarity effect previously observed by Vinckier et al. (2007) was due to lower resolution data, image smoothing, and inter-subject averaging.

Second, we evaluated the hypothesis that a succession of patches along the posterior to anterior axis of the VOTC may be selectively sensitive to increasingly complex orthographic components. According to this hypothesis, patches would be specialized for increasingly larger components of words, from frequent letters, to bigrams, quadrigrams, culminating in anterior regions sensitive to lexical status (Dehaene et al., 2005). Following this hypothesis, the word similarity effect we observed could actually be due to discrete steps of activity increase, with each patch responding to an orthographic unit of a certain grain size. Alternatively, the activity increase of the word similarity effect may be genuinely continuous in nature: as stimuli get increasingly similar to words in the orthographic lexicon, they would generate a bottom-up activation which increases monotonously, or perhaps they could also receive progressively increasing top-down feedback signals from higher areas (Woolnough et al., 2021). In order to test these hypotheses, at this stage irrespective of language, we contrasted conditions in which the frequency of orthographic components was high versus low in both languages (L+ versus L-, B+ versus B-, and Q+ versus Q-), and contrasting words (WE, WF) versus pseudowords (QE, QF) for the lexicality effect.

We first examined the prevalence of whole-brain activation across individual participants. Lexicality elicited relatively consistent differences between words and pseudowords in 76% (16/21) of participants, with clusters of higher activation for words in 15 participants, and clusters of higher activation for pseudowords in 7 participants (p<0.001 uncorrected, cluster size > 4). Higher activation for words (number of participants in brackets) was mainly found in language areas along the STS (13), in IFS/IFG (8), precentral sulcus (5), supramarginal gyrus/planum temporale (5), MFG (3), SMA/preSMA (3), IPS (3). Only four participants showed positive activation in the VOTC, and in 3 of them activation was located around the middle occipito-temporal gyrus and superior to the VWFA. Conversely, higher activation for pseudowords was mainly found along the brain midline and close to the frontal pole, including the retrosplenial cortex (3), parieto-occipital junction (3), precuneus (2), medial superior frontal gyrus (2), MFG (2). None of the other planned contrasts elicited any activation in more than 5 participants, and the consistency across participants was low. In summary, while lexicality effects (word versus pseudoword) induced discrete activity jumps, other levels (letters, bigrams, or quadrigrams) did not.

We therefore moved to analyses within the 773 word-specific ROIs obtained from the localizer analysis. For each ROI, we averaged time courses across voxels, and re-computed the GLM and contrasts (ROI-GLM). We examined the same contrasts as in whole-brain analyses (**Figure 4**, top row), plus the pairwise differences in frequency and lexicality effects between successive component types (B vs L, Q vs B, W vs Q, **Figure 4**, bottom row). Overall we only found a very small number of ROIs sensitive to frequency for each component level (L: none; B: 7; Q: 23), only few of which were in the VOTC (L and B: none; Q: 7). The lexicality effect (W > Q) again showed more robust activation in lateral temporal and frontal areas (ROIs from 9 participants, including 6 SB, 2 SF, 1 SE), but again with few VOTC ROIs (4 ROIs, from 2 SB). This scarcity of significant ROIs is consistent with whole-brain analyses, and indicates that reading-related patches are not selectively sensitive to discrete, specific levels of orthographic structure, but are gradually activated by increasingly word-like stimuli. Consistent with this conclusion, note that the vast majority of ROIs already showed above-baseline activity even for the infrequent letter condition (593/773 ROIs across the brain and 285/299 VOTC ROIs at FDR q<0.05). While it remains possible that successive word patches respond to other increasingly invariant features of reading such as size, case or font invariance (Dehaene et al., 2004), as found for face recognition (Freiwald & Tsao, 2010; Tsao et al., 2008), their organization does not separate neatly according to the exclusive presence of letters, bigram or quadrigrams.

**Figure 4.**
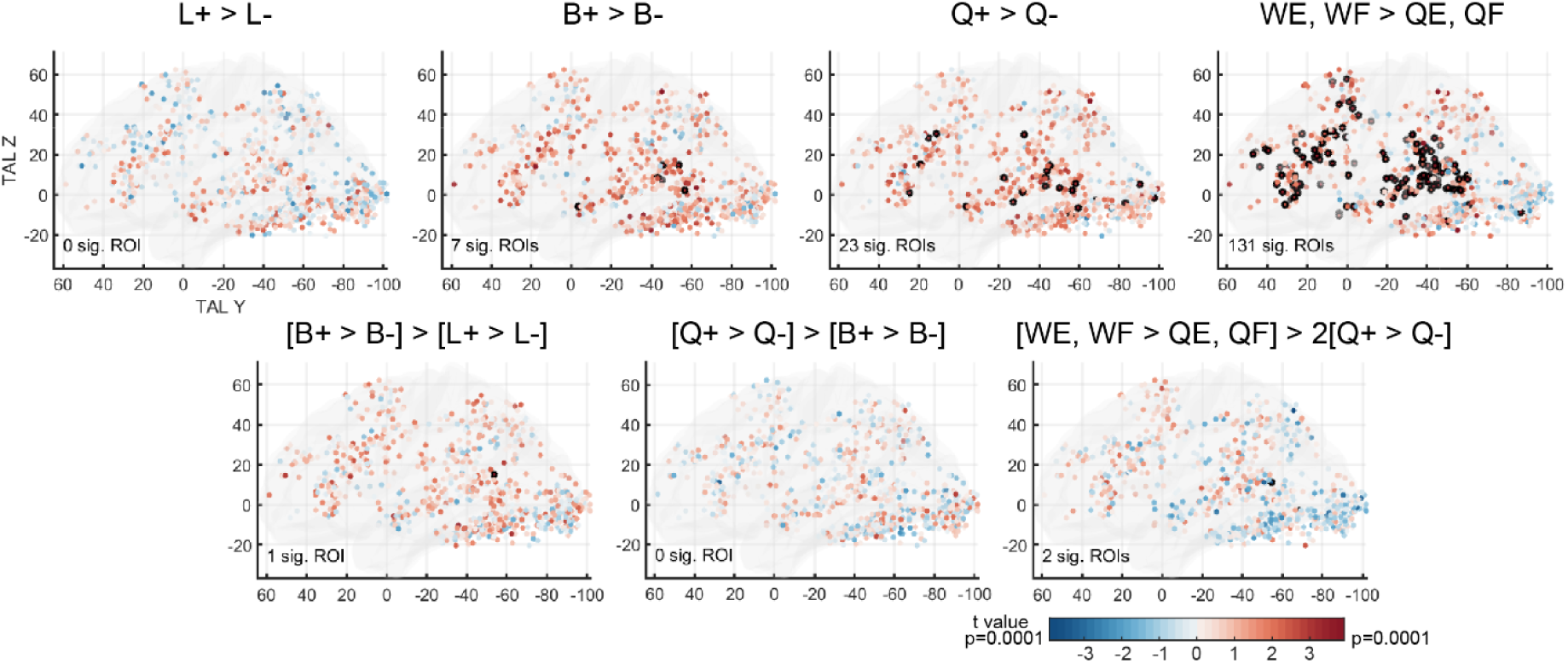
Contribution of letter, bigram and quadrigram frequencies, as well as lexicality, to the overall word similarity effect in English-French bilingual readers. Top row: Contrasts of high minus low frequency conditions for letters, bigrams, and quadrigrams, and contrast of real words minus matched quadrigrams, in the 773 word-specific ROIs. Bottom row: pairwise comparisons between the contrasts depicted in the top row. ROIs circled in black are significant after correction for multiple comparisons. (FDR q<0.05, corrected within the VOTC, lateral temporal, or lateral frontal region, respectively. No FDR correction for the 107 ROIs falling outside these 3 regions.) ROIs in the VOTC showed no effect, providing no evidence that they are specialized for a specific type of orthographic components. Lateral temporal and frontal ROIs were mainly sensitive to lexical status and quadrigram frequency.

#### Do distinct cortical patches specialize for English versus French?

The main goal of our experiment was to search for putative cortical patches with a specific tuning to English or French. To this end, lexicality and frequency statistics were manipulated orthogonally between the two languages.

### Differences between English and French real words

The localizer experiment revealed no consistent differences between activations to English and French words. This conclusion was confirmed in the main fMRI runs, despite having much more data than in the localizer (15 versus 5 repetitions per condition). Only 4 participants showed seemingly language-specific activations in a total of 29 clusters, but those findings were not consistently replicated in the localizer: Within the 29 main-experiment clusters, we tested the differences between English and French in the localizer data with ROI-GLM. The effect of language was non-significant (22/29 ROIs) or weak (0.005<p<0.05, 6/29 ROIs). Similarly, in the 773 word-specific ROIs, ROI-GLMs identified only 6 ROIs with a significant language difference (FDR q<0.05).

### Differences between English and French sublexical statistics

Within word-specific ROIs, for each sublexical component (letter, bigram, quadrigram), we compared stimuli whose component frequency diverged between English and French (LE vs LF, BE vs BF, QE vs QF). Only a single ROI showed a language difference for bigram frequency. We also assessed the full 2 × 2 design by probing (1) the main effect of frequency separately for each language (e.g. [LF, L+] vs [LE, L-] for the main effect of letter frequency in French), and (2) the interaction term, which would indicate that the frequency effect was significantly larger in one language than in the other. Crucially, this interaction term was never significant for any of the sublexical components. We only found two ROIs with a significant main effect of bigram frequency in English, and a few ROIs with a significant main effect of bigram or quadrigram frequency in French (34 and 12 ROIs, respectively). These ROIs came from 7 participants (6 SB, 1 SF), although the majority of ROIs actually came from one French participant (SF01, bigram effect: 20 ROIs, quadrigram effect: 5 ROIs).

### Effects of individual language dominance

Our 21 participants comprised 7 English-dominant, 7 French-dominant, and 7 balanced bilinguals. We examined whether their language profiles were reflected in the brain, namely the activity difference between English and French conditions across individual participants. For each participant, we computed a behavioral score of language dominance based on the number of words they read aloud in one minute: ([English words - French words]/[English words + French words]). To obtain a comparable index at the brain level, we merged bilateral word-specific ROIs into bigger anatomical regions or sub-regions for each participant. The 3 broad anatomical regions were the same as defined before (VOTC, lateral temporal, lateral frontal, see **Figure 3A**). For a detailed examination of VOTC and especially around the fusiform, we further examined sub-regions of ROIs surrounding the fusiform gyrus according to specific individual anatomical locations, including the mid-fusiform sulcus (mFS) and the IOS-OTS sub-regions; and we grouped all VOTC ROIs posterior to the fusiform region into an early visual cortex (EVC) region. Within each region, we computed the average activity difference evoked by English vs French conditions in the localizer and main runs (WE-WF, LE-LF, BE-BF, QE-QF). Finally, we correlated the activity difference with each participant’s behavioral language dominance score.

For activity differences between English and French words of the main fMRI runs, as expected, no significant correlation was found in EVC (r(19)=-0.183, p=0.427). In the fusiform region, however, a negative correlation was present (r(19)=-0.550, p=0.0098, FDR corrected for 5 comparisons within each region, **Figure S5**), indicating that words in the dominant language induced a *lower* activity, presumably due to lower effort. Dissecting the fusiform region, this negative correlation arose from the IOS-OTS sub-region (r(19)=-0.661, p=0.0011), but not the mFS (r(15)=0.357, p=0.159). Outside the fusiform region, a similar negative correlation was also found in lateral temporal and lateral frontal regions, although not surviving FDR correction (r(19)=-0.548 and r(14)=-0.600; uncorrected p=0.0102 and p=0.014, respectively). None of the other sublexical components (LE-LF, BE-BF, QE-QF) correlated with the behavioral language dominance in any of the regions/sub-regions (all p > 0.064). The language difference in the localizer did not show any significant correlation either (all p > 0.339).

#### Summary of the English-French experiment

With the higher resolution afforded by 7T compared to previous 3T studies, the VWFA became subdivided into a multitude of cortical patches that were highly word-specific, and required single-subject analyses. Activity showed a robust word similarity effect in 64.5% of word-specific ROIs, and a posterior-to-anterior increase of this effect along the VOTC, reproducible in every single participant. However, all of those ROIs were jointly co-activated by English and by French words, and we did not find consistent differences between languages, either with whole-brain or with ROI analyses.

The lack of language difference in the VOTC of English-French participants may be due to the fact that English and French use identical alphabets. To examine whether widely different scripts lead to activity differences in the VOTC, we extended our experiment to 10 English-Chinese bilingual readers.

### Experiment 2: Reading in English-Chinese bilinguals

The experimental design was very similar to the English-French experiment (**Figure 5**). The localizer used the same design, except that French words and checkerboards were replaced with Chinese words and with scrambled strokes derived from these words, respectively. The main fMRI runs comprised two distinct hierarchies of English and Chinese stimuli with increasing similarity to real words. The English stimuli were directly taken from the English-French main fMRI runs, i.e. the conditions in which the frequencies of orthographic components were congruent in English and French (L-, L+, B-, B+, Q-, Q+) plus the real English words (WE). Chinese stimuli included strokes covering the entire area where two Chinese characters could appear; strokes organized in 2 groups similar to two Chinese characters; Chinese radicals arranged to form 2 pseudo-characters (with radicals arranged in either orthographically impossible or possible positions); real Chinese characters whose pairings formed non-words; and finally real Chinese words, 2 characters long, of low and high frequency (**Figure 5**, see Materials and Methods for details).

**Figure 5.**
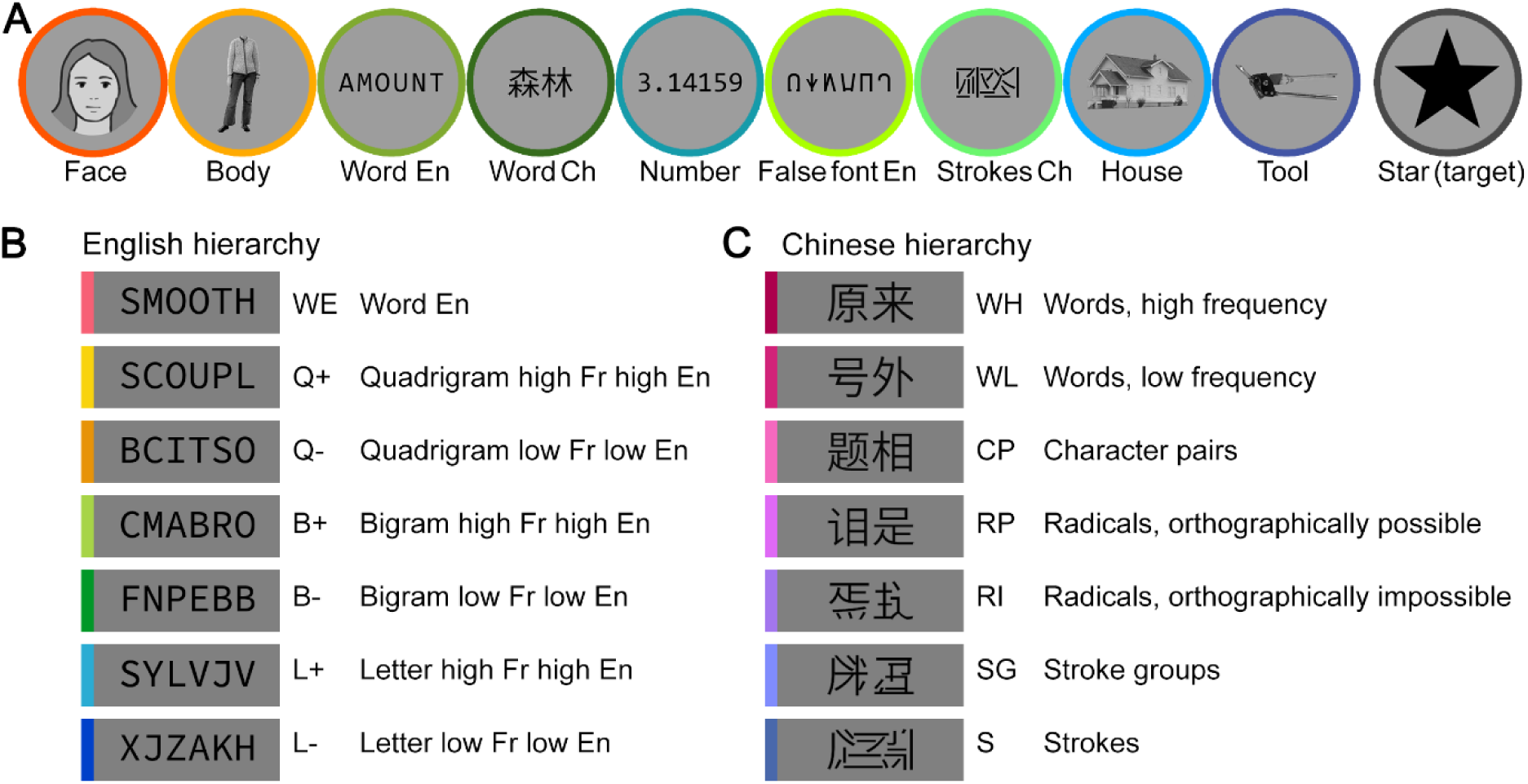
Stimuli and procedure for experiment 2 with English-Chinese bilingual readers. **A.** Examples of the 9 categories of visual stimuli, plus the target, used in the localizer. Most of the conditions were the same as in the English-French experiment, with Word Ch replacing Word Fr, and Strokes Ch replacing Checkerboards. The actual face stimulus is replaced by an illustration here. **B. and C.** Design of the word similarity experiment. **B.** English stimuli, identical to the corresponding conditions in the English-French experiment (see **Figure 1**). **C**. Chinese stimuli with hierarchical components increasingly similar to real words (see Materials and Methods for details).

#### Localizer for visual categories and overall reading circuit

For consistency, we performed the same bilingual word-specific contrasts as in the English-French study, irrespective of languages (English and Chinese words > faces, bodies, houses, tools). Similar to the English-French study, the localizer revealed that every individual English-Chinese participant possessed robust word-specific activation clusters in the VOTC, lateral temporal, and lateral frontal regions (296 ROIs across participants, **Figures 6A-B and S6**). In the VOTC, the mean number of word-specific clusters and voxels was again larger in the left hemisphere (paired t-tests, 7.9 vs 2.1 clusters, p=0.0079; 496.2 vs 71.7 voxels, p=2.99×10^-4^; no right-hemisphere voxel for participants SC05 and SC10). However, the bilingual word-specific contrast overlapped significantly less with voxels from the single language-specific contrasts (whole brain, English: mean=68.01%, SD=25.39%; Chinese: mean=71.70%, SD=16.12%; VOTC, English: mean=64.40%, SD=22.51%; Chinese: mean=79.90%, SD=13.04%), compared to the high overlap in English-French participants (Wilcoxon rank sum test, whole-brain: p=4.898×10^-4^, VOTC: p=1.482×10^-5^). This indicates that the bilingual word-specific contrast in the localizer is likely missing language-specific voxels, and the two languages are potentially more segregated in the English-Chinese participants.

**Figure 6.**
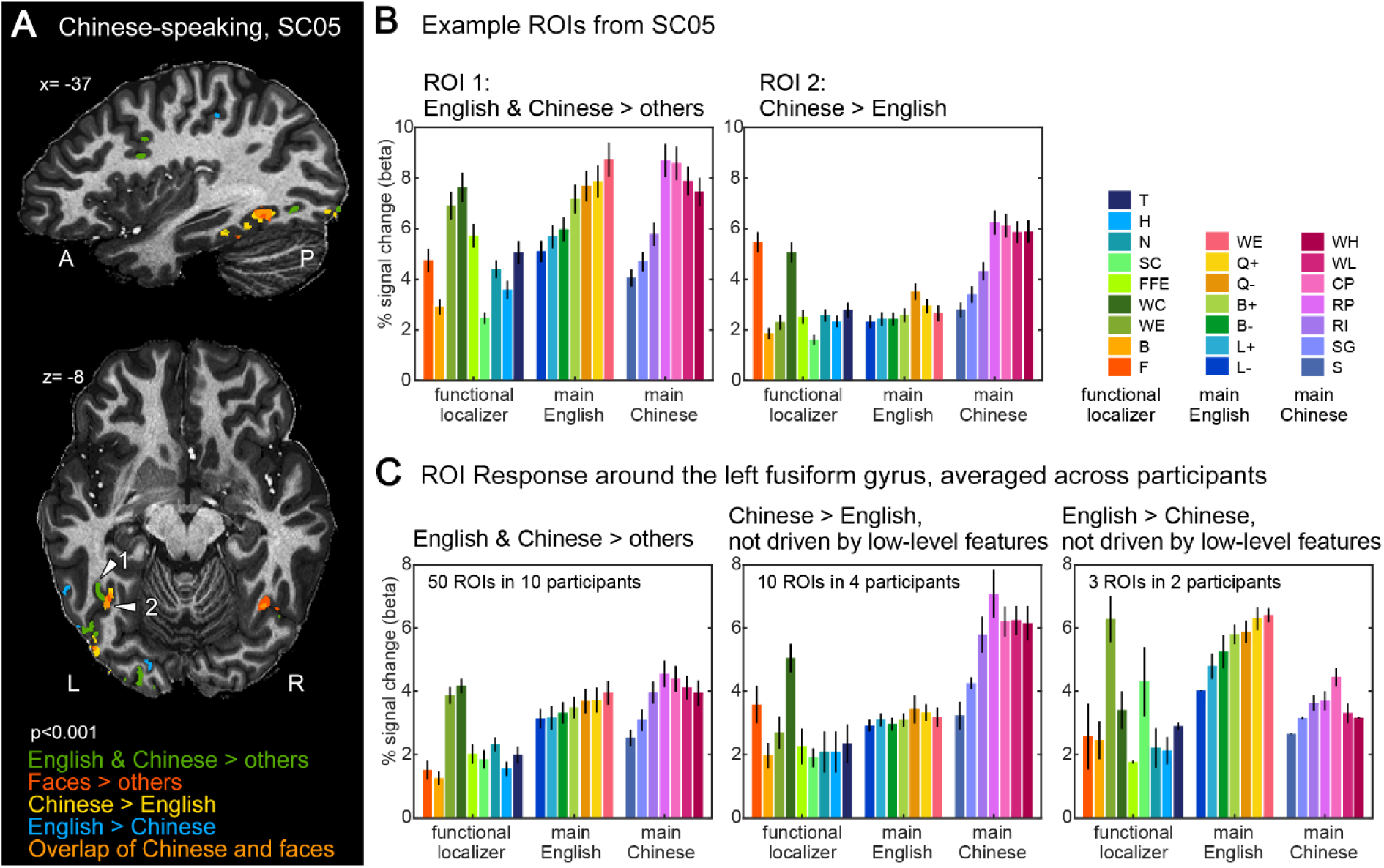
Word-specific and language-specific activation in English-Chinese bilingual readers. **A.** Example of ventral occipito-temporal activation in a representative participant (SC05, p<0.001 uncorrected. See **Figure S6** for the maps of all 10 participants). The word-specific and face-specific clusters were defined with the localizer (English and Chinese words > faces, bodies, houses, tools; faces > bodies, houses, tools, English and Chinese words). The Chinese- and English-specific clusters were defined with the main fMRI runs (Chinese words > English words and vice-versa). Note the overlapping sensitivity for Chinese words and for faces (orange color) in ROI 2 (white arrow) and a symmetrical right-hemispheric ROI. **B.** Activity profile of two example ROIs from participant SC05, labelled with white arrows in panel A. ROI 1 shows shared responses to English and French words and corresponding word similarity effects, while ROI 2 is sensitive to Chinese stimuli as well as faces. **C.** Group-average activity profiles of bilingual word-specific ROIs around the left fusiform gyrus (left), with preference for Chinese stimuli (middle) or with preference for English stimuli (right). In the Chinese-specific ROIs (middle panel), note the strong response to faces (red bar). See **Figure S7, S8** for all individual ROIs.

We also found face-specific clusters in bilateral VOTC of every participant, although the numbers of clusters and voxels were not significantly different between left and right hemispheres (5.2 vs 5.8 clusters, p=0.452; 264.8 vs 352.7 voxels, p=0.252).

#### Spatial gradient of word similarity

Within these bilingual word-specific clusters, we assessed the word similarity slopes and gradients similar to the experiment on English-French bilinguals, but now separately for English and Chinese (**Figure 7**, comparable to **Figure 3A**). We did not perform FDR correction here, due to the much smaller number of conditions being fitted (first 5 conditions per language in Experiment 2, versus all 14 conditions in Experiment 1. See Materials and Methods). Within the 296 ROIs, 74 and 40 showed a significant slope for Chinese and English, respectively, including 23 and 15 ROIs in the VOTC (p<0.05, uncorrected). In the left VOTC, there was a significant increase of the word similarity effect along the posterior-to-anterior axis for both English and Chinese (word similarity slopes fitted with TAL Y coordinates in individual participants, one-sample t test for the resulting regression coefficients against 0, English: t(9)=2.755, p=0.0223; Chinese: t(9)=4.161, p=0.0024). Note that since the English and Chinese conditions did not have one-to-one correspondence, the word-similarity slopes and their spatial gradients cannot be meaningfully compared between languages.

**Figure 7.**
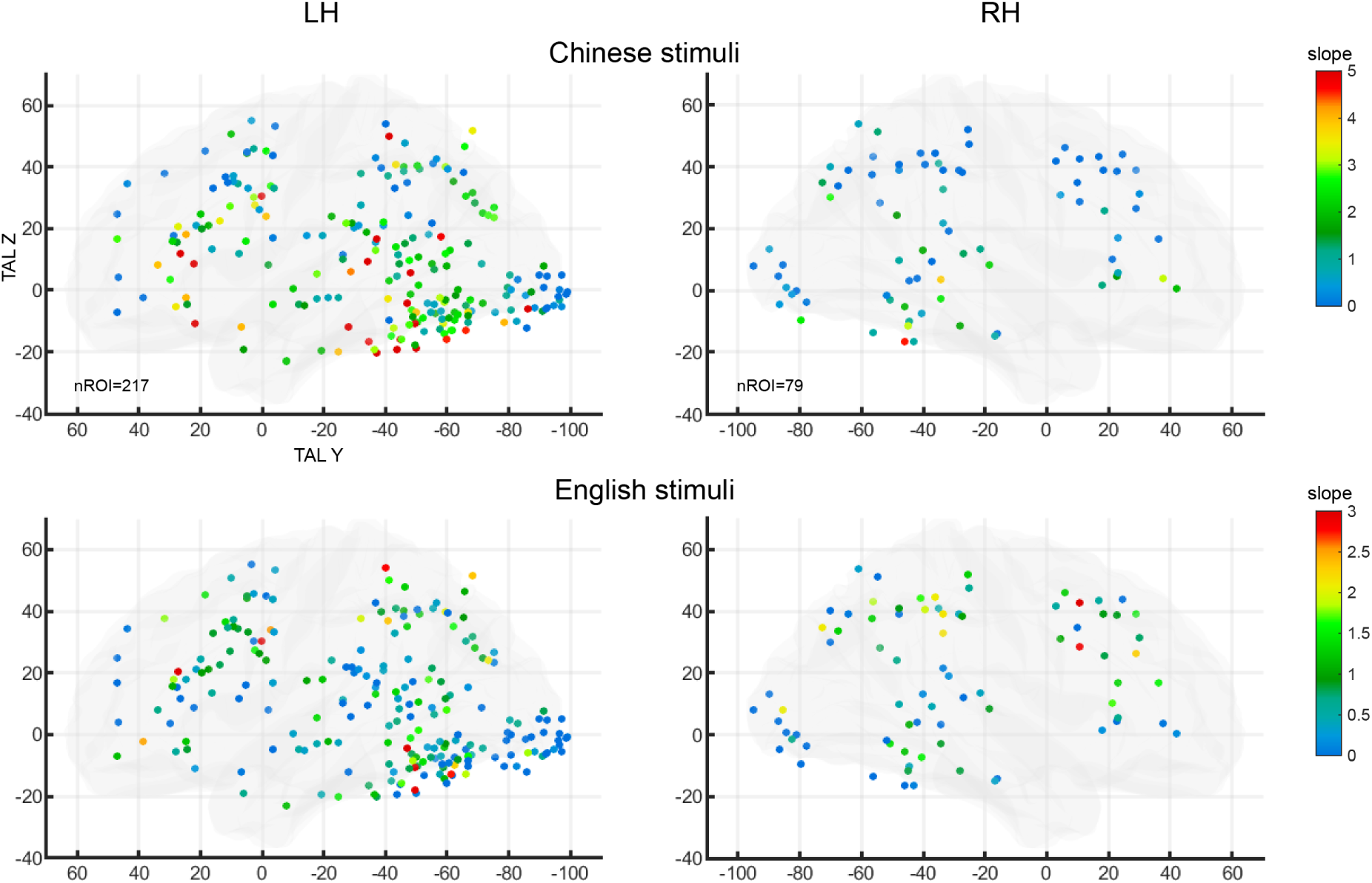
The word similarity effect in word-specific ROIs of English-Chinese bilingual readers. Same format as **figure 3A**. Word similarity was modelled as the slope of a linear regression over the first 5 stimulus conditions for each language. Slopes for Chinese and English word similarity effects are color-coded for each of the 296 ROIs from the contrast English & Chinese > others in the localizer. Note the scales for Chinese (top) and English stimuli (bottom) are different.

As in the English-French study, we assessed the hypothesis that discrete cortical patches would be sensitive to different hierarchical levels of stimulus structure. We performed ROI-GLM contrasts in the word-specific ROIs, comparing pairwise conditions. For English stimuli, as in experiment 1, we found no effect of frequency for letters (L+>L-), bigrams (B+>B-) or quadrigrams (Q+>Q-), but an effect of lexicality effect in lateral temporal and lateral frontal regions. For Chinese stimuli, the pairwise contrasts between the 4 consecutive non-character conditions (SG>S, RI>SG, RP>RI) each showed significant ROIs spread along the VOTC, with no evidence for the processing of hierarchic levels being associated to distinct anatomical regions. When contrasting real characters with orthographically possible radicals (CP>RP), significant clusters were mostly located in the lateral temporal and lateral frontal areas, similar to the lexicality effect for English stimuli. Finally, other contrasts involving real characters (WL>CP, WH>WL) did not show any significant ROI.

We analyzed the effects of language dominance as in experiment 1, but did not find any significant correlations in any brain regions or sub-regions (all p > 0.084). This may be because the sample size of English-Chinese bilinguals are smaller (n=10), and all of them were dominant in Chinese.

#### Cortical patches specialized for Chinese

Different from Experiment 1 with English-French bilinguals, in English-Chinese bilinguals the localizer data is likely missing some language-specific clusters, since the bilingual word-specific contrast overlapped much less with the single-language-specific contrasts. To maximize sensitivity and avoid false negatives, we localize them using the data from the main fMRI runs (more data, 15 repetitions versus 5 in the localizer), and then replicated and extended the findings using the independent data from the localizer.

These analyses revealed robust Chinese-specific clusters in the majority of participants. Contrasting Chinese words (high, low frequency) > English words (p<0.001 uncorrected, cluster size > 4) resulted in 66 bilateral activation clusters, mostly located around the fusiform gyrus (e.g. in OTS and IOS, **Figure S6**; 31 clusters, 8/10 participants), and in the STS (24 clusters, 5/10 participants). The fusiform clusters were more numerous in the left than in the right hemisphere (mean 2.87 vs 1.00 clusters, p=0.0058), which did not differ from the leftward bias observed with the bilingual English-Chinese clusters (χ^2^(1)= 0.758, p=0.3840). The remaining 11 clusters were from a few participants, including 5 early visual area clusters from 3 participants (SC05, SC08, SC10), 5 frontal clusters from two participants (SC04, SC06), and 1 IPS cluster from SC10. In most clusters, the Chinese specificity was replicated with independent data from the localizer (ROI-GLM contrasts on the localizer time courses): the contrast of Chinese > English words was significant in most clusters across participants (p<=0.001 in 31/66 clusters, 0.05>p>0.001 in another 24/66 clusters, no FDR correction) and failed to reach significance in only 11/66 clusters. As for the Chinese-specific clusters located around the fusiform gyrus (in OTS/IOS/IOG), the language preference for 87% (27/31) of those clusters was systematically replicated in the localizer (all p<0.05, FDR q<0.05).

Focusing on these 31 fusiform Chinese-preferring clusters, we further examined whether the language preference could be explained by low-level retinotopic differences between the stimuli. If this was the case, the same difference between languages should be present between the matched low-level control conditions. Hence we assessed the interaction (English > Chinese words) - (English false fonts > Chinese strokes) in the localizer. The interaction reached significance in 11/31 clusters (10 of which in the left hemisphere from 4 participants), demonstrating Chinese specificity independent of low-level features (see **Figure 6C** for averaged activity profiles across participants, and **Figure S7** for individual clusters; the latter figure shows that in fusiform/OTS clusters, even when the interaction failed to reach significance, its direction was almost always consistent with a genuine language-specific effect).

The fusiform clusters showed further signatures of selective tuning to the Chinese script in the word similarity slopes. The Chinese-specific clusters showed a significant word-similarity slope only across Chinese conditions, but not across English conditions (p<0.05 in 12/31 ROIs for Chinese, 0/31 ROIs for English, no FDR correction in order to avoid false negatives). Visual inspection confirmed that most fusiform ROIs had a distinctive activity profile characterized as: (1) a nearly flat activity profile across the 7 conditions with variable similarity to English words; (2) an increasing activity across the first 4 conditions with variable similarity to Chinese words; (3) a high activity for the last 4 conditions, i.e. for any orthographically possible combinations of Chinese radicals, with a trend for a decrease in activity for real characters and words. This finding indicates that those regions, much like the classical VWFA, are prelexical in nature and can be strongly activated by pseudowords with the right kind of subunits.

The localizer also led to a surprising finding of overlap between face-specific and Chinese-specific activations (**Figure 6A**). While Chinese-specific VOTC clusters showed a much higher activity to Chinese words than to almost any other categories in the localizer, including English words, there was one exception: they were strongly activated by faces (**Figure 6**, **Figure S7**). 25/31 ROIs showed significant face specificity in the ROI-GLM (faces>bodies, houses, tools, 14 ROIs with p<=0.001; 11 ROIs with 0.001<p<0.05). Such specificity for faces remained true in the subset of clusters whose Chinese specificity was demonstrably not driven by low-level features (**Figure 6C central panel**). Furthermore, Chinese-specific clusters were often overlapping with or very close to face-specific clusters (**Figure S6**, yellow arrows).

#### Very few cortical patches specialized for English

In the opposite direction, for the contrast of English > Chinese words, one participant did not have a single English-specific cluster (SC06) and another participant had an abnormally large number of them (SC03: more than 200 clusters at p<0.001, 46 clusters even at a more stringent threshold of p<0.0001, mainly in frontal, early visual area and middle occipital gyrus). Leaving aside those two outliers, the majority of the clusters for this English-specific contrast for the rest of 8 participants were consistently found in early visual areas (23/62 clusters) and were due to visual rather than linguistic preference, since 22/23 did not pass the criterion of showing critical interaction relative to lower-level stimuli (Chinese > English words) - (Chinese strokes > English false fonts). Other clusters were found in the OTS/IOS region (11/62 clusters in 5 participants; **Figure S8**), and 5/11 of them passed the critical interaction for English specificity not driven by low-level differences (**Figure S8A, C**, 3 clusters in the left hemisphere). These 5 clusters therefore qualified for English specificity, but 3 of them showed a significant word-similarity slope not only for English but also for Chinese (p<0.05, no FDR correction). The remaining 28/62 English-preferring clusters came from only 3 participants and were mainly located in the IPS and lateral frontal areas.

In summary, as compared to Chinese-specific activations, English-specific clusters were few, even fewer in the VOTC, and their language-specificity was weak.

### Data-driven comparison of English-French and English-Chinese bilinguals

Looking for evidence of language specialization in the VOTC requires pushing spatial resolution as much as possible, which is why we used 7T imaging at 1.2 mm isotropic. Still, each of our voxels comprised ∼150-200 thousand neurons with different functional properties – and furthermore, many of the above analyses, to achieve adequate signal-to-noise, averaged the signal over multiple voxels in a cluster that we implicitly supposed to be homogeneous. In a final analysis, we try to side-step this limitation by using “hypothesis-free decomposition” (S. Norman-Haignere et al., 2015), a non-parametric ICA which can infer the multiple canonical response profiles of distinct neural populations possibly overlapping within the same voxel. This method was previously shown to identify distinct neural responses to music and to speech in 3T fMRI (S. Norman-Haignere et al., 2015), which were later replicated and extended by direct intracranial analysis (S. V. Norman-Haignere et al., 2022).

We used this approach to decompose the response profiles from the main fMRI runs, within all the bilingual word-specific VOTC voxels defined by the localizer data (experiment 1: English & French words > others; experiment 2: English & Chinese words > others, p<0.001 uncorrected, cluster size > 4). For each group of bilinguals, we decomposed the 14 conditions into 3 canonical response profiles, i.e. the minimum number required to model baseline activity, the word similarity effect, and putative language differences. Activity in each voxel was expressed as a weighted sum of those 3 profiles, each multiplied by a voxel-specific weight.

Figure 8 shows the data-driven component profiles that were identified by this procedure. Although the components were identified solely from the 14 rightmost conditions (main fMRI runs only), for completeness the figure also shows the corresponding activity profiles from the localizer run, which were obtained by applying to each voxel the component weights from the main fMRI runs. In both language groups, we observed a flat component showing similar activity amplitude across all conditions, similar to activity in early visual cortex (compare **Figures 8** and **3B**). The other components confirmed that English-French and English-Chinese bilinguals differed sharply. For English-French participants, the second component showed an overall word-similarity effect, which was nearly perfectly continuous and monotonic, with no differences between languages. The third component was noisy, and exhibited an overall quadratic organization, presumably capturing between-voxel variations in the shape of the word-similarity effect. Although it did show hints of language differences (higher amplitude for LF, BF, QF, WF than for LE, BE, QE, WE, respectively), this pattern was not consistent with the localizer profile, where WE was higher than WF, i.e. in the opposite direction.

The results were strikingly different in English-Chinese participants (**Figure 8**, bottom row). The word-similarity effect was dissected into two distinct components, one for English (component 3) and one for Chinese (component 2). Each showed a higher amplitude for stimuli in the preferred language, whether words or pseudowords. Furthermore, this language difference was replicated in the localizer profiles. Importantly, these language differences were not driven solely by the Chinese- and English-specific voxels. First, only a small proportion (404 and 257 out of 5679 voxels, 11% in total) of the word-specific voxels analyzed here (as defined by the localizer data) overlapped with the Chinese- or English-specific voxels (as defined by the main fMRI runs). Second, removing those language-specific voxels did not qualitatively change the shape of the component profiles.

**Figure 8.**
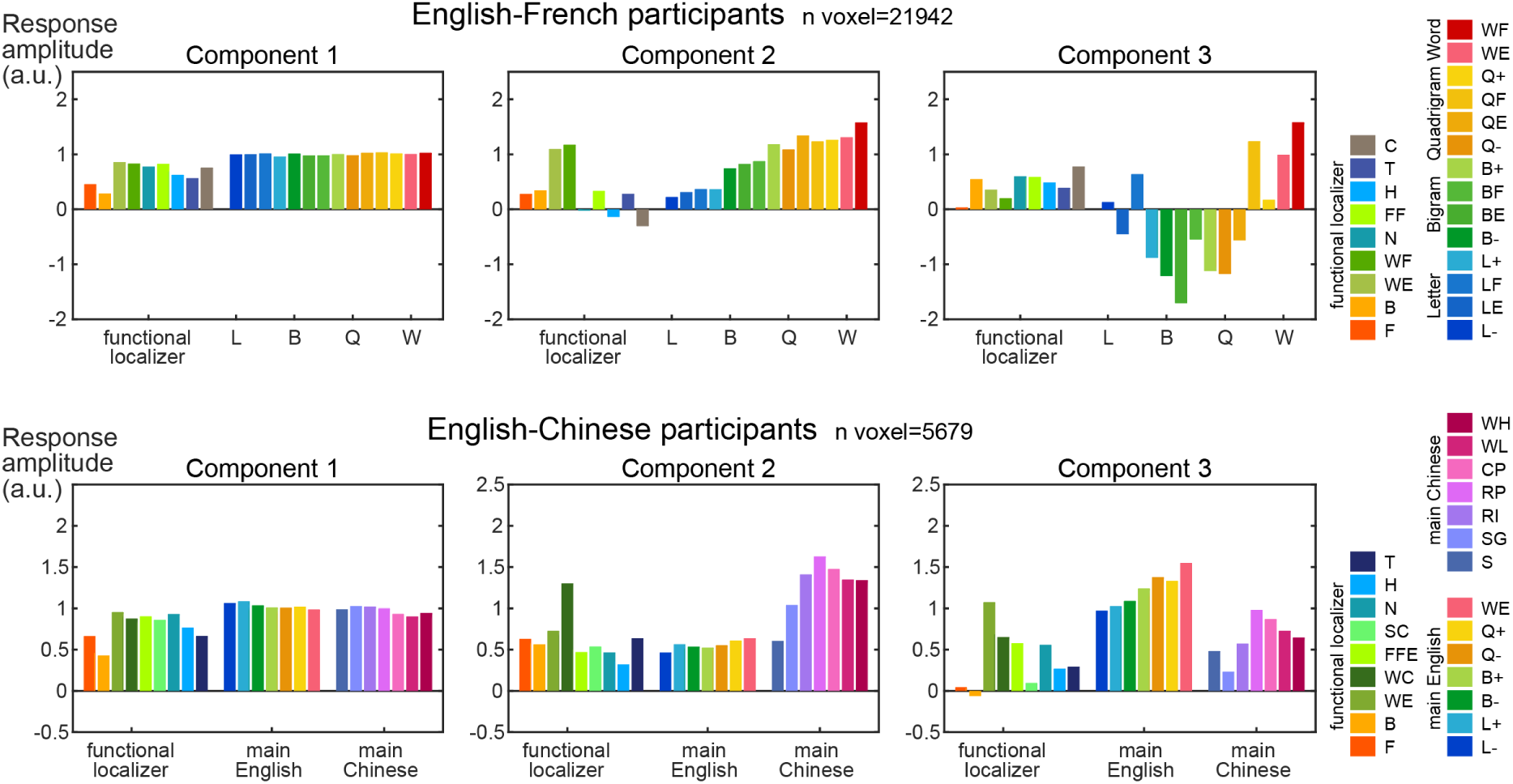
Data-driven decomposition of activity profiles in ventral occipito-temporal cortex. A variant of independent-component analysis (ICA) was applied to the activity profiles of the main fMRI runs (14 rightmost bars in each graph), within each word-specific voxel identified in the localizer. The resulting weights were also applied to the localizer data to derive their activity profile (leftmost bars). In English-French participants, activity was decomposed into a weighted sum of constant, linear and non-linear shapes of the word similarity effect, without any clear separation between English and French stimuli. In English-Chinese participants, by contrast, components 2 and 3 clearly showed distinct preferences for either Chinese or English stimuli.

In summary, those analyses demonstrated the added value of a purely data-driven ICA analysis: this analysis managed to identify sub-voxel functional properties, and confirmed the presence of a language specialization in English-Chinese, but not in English-French participants.

## General discussion

In the current 7T fMRI study, we examined the organization of reading-related cortical patches in English-French and English-Chinese bilinguals. Because we operated at a high resolution and SNR, our analyses were entirely focused on individual participants. In spite of inter-individual variability, three general conclusions could be drawn: (1) the VWFA actually consists of multiple cortical patches with high selectivity for written words; (2) identical patches respond to English and French stimuli in bilingual English-French readers, but partially distinct patches respond to Chinese and English scripts in bilingual English-Chinese readers; (3) patches are highly sensitive to language statistics. We discuss these points in turn.

### Multiple specialized cortical patches for reading

By analyzing unsmoothed fMRI data at 1.2 mm isotropic resolution, we found that what previously appeared, at a lower resolution, as a single extended visual word form area (VWFA) actually consists in a multiplicity of small and highly specialized word-specific patches spread in the VOTC along the OTS/IOS. These reading-related patches bear similarity to the face patches found both in the current study and in previous literature, although showing perhaps less stereotypical distribution across participants (Arcaro et al., 2020; Tsao et al., 2008). For both English-French and English-Chinese participants, word patches exist in both hemispheres, but with a left-hemisphere bias in numbers of clusters and voxels.

We largely prevented the usual signal drop in the anterior inferior temporal lobe (between TAL Y=-50 and -22), and found that this region also contains cortical patches with high word selectivity. These clusters may correspond to the supramodal “basal temporal language area” in the intracranial recording literature (Cohen et al., 2021; Krauss et al., 1996; Schaffler et al., 1996). In one recent intracranial study, the word-selective sites anterior to TAL Y=-40 were more responsive to words than false fonts and infrequent letters under a passive viewing task (Woolnough et al., 2021), consistent to our results (**Figure S4**).

Given the numerous word patches found along the OTS/IOS, and the difficulty to arrange them in larger groups reproducible across participants, the word patches may be better studied individually. By analyzing them one by one, we identified several local cortical properties. First, word patches can be exquisitely specialized, showing much lower activity for non-word stimuli with similar low-level features, such as numbers, false fonts and character strokes, and also other object categories with less similar visual features (bodies, houses, tools). In this respect, 7T fMRI converges with intracranial recordings in suggesting that the VOTC is a patchwork of small, highly specific cortical sectors each responsive to a small category of stimuli, including learned ones such as letter or number strings (McCarthy, 1999; Nobre et al., 1994; Puce et al., 1996). An analogy with face patches suggests that word patches may contain a vast majority of neurons highly specialized for letters and written words (Tsao et al., 2006) – a prediction that may become testable as high-density single-cell recordings become available in humans (Chung et al., 2022; Paulk et al., 2022). Supporting this prediction, simulations of neural networks designed for object and face recognition but recycled for written word recognition show that a small proportion of units becomes entirely dedicated to reading (Hannagan et al., 2021).

However, a surprising finding was that many of the Chinese-specific patches also showed high face specificity (faces>bodies, houses, tools) (for related findings, see J. Liu et al., 2008, 2009). Chinese characters may share with faces a need for holistic (or configural) processing (Farah et al., 1998; Richler & Gauthier, 2014, p.). While alphabetic writing also calls for encoding the relative position of letters (Grainger & Whitney, 2004; Jordan et al., 1999; Wong et al., 2011), information may be more configural in Chinese characters (Chen et al., 2013; Mo et al., 2015; Tso et al., 2021; Wong et al., 2012), which feature symmetries, horizontal and vertical alignments, and other regularities in two dimensions. Additionally, the shared overall round shapes of Chinese characters and faces may be preferred by a ventral visual network for “stubby” shapes, as reported in monkeys (Bao et al., 2020).

The cortical overlap between Chinese characters and faces is reminiscent of the frequent interactions that have been observed between reading and face recognition during the acquisition of literacy. When comparing literate and illiterate participants, the activity evoked by faces tended to decrease with literacy in the left-hemisphere VWFA location and showed a strong increase in the right hemisphere (Dehaene et al., 2010). This previous finding suggested a competition between the newly acquired words and the already existing face specificity, consistent with the cultural recycling hypothesis (Dehaene & Cohen, 2007). The word-face competition hypothesis was revisited in later developmental studies of children learning to read (Dehaene-Lambertz et al., 2018; Feng et al., 2022; Monzalvo et al., 2012): during reading acquisition, the VWFA invades largely unspecialized cortical patches in the left OTS, and their growth blocks the slow development of face patches and forces it to shift towards the right hemisphere (Feng et al., 2022; Golarai et al., 2010), perhaps because both call upon the same cytoarchitectonic region (Gomez et al., 2017). The current study confirms that, indeed, there may be shared mechanisms between face and word processing, but surprisingly indicate that cortical competition (in alphabetic readers) may switch to cortical overlap (in Chinese readers). Whether this overlap is reflected in a performance enhancement (potential skill transfer) should be clarified in future studies.

### Script-specific patches

The main goal of our study was to ask whether the VWFA splits in bilingual readers. We found that the answer depends on whether the languages share the same script or not. For English versus Chinese, we found a partial split: several VOTC clusters were highly specific for Chinese only and showed the word similarity effect only for Chinese, while very few showed the converse preference for English over Chinese. Even within voxels selected for their reading-related activity regardless of language, a data-driven analysis indicated the presence of subvoxel preferences for one or the other language (**Figure 8**). There was no such separation, however, when English and French stimuli were contrasted in English-French bilinguals. This absence of a split is unlikely to be due to lack of power, because we had 21 participant in the English-French group, twice the size of the English-Chinese group. The same word similarity slope for both English and French conditions could be observed in individual participants, ROIs, and even single 1.2 mm voxels, without showing any consistent language bias. In reading-related clusters around the fusiform area, we even detected a subtle dominance effect across participants, where the activity balance between English and French words was in favor of the less dominant language (**Figure S5**), which we interpret as greater top-down attention or cognitive effort. In the sub-voxel profiles, the analyzed voxels for the English-French group were also almost 4 times as numerous as for the English-Chinese group – and yet no language split was detected.

The discovery of Chinese-specific voxels was made possible here thanks to the small voxel size (1.2 mm isotropic) and high signal-to-noise afforded by 7T fMRI. Using coarser resolution, prior studies mostly failed to find differences between the cortical circuits for reading, both between languages using the same alphabet, and between Chinese/Japanese and alphabetical languages (Rueckl et al., 2015). This is notably true for VOTC activations both in monolingual and bilingual individuals (Brignoni-Perez et al., 2020; Hsu et al., 2015; Jamal et al., 2012; Li et al., 2021; Meschyan & Hernandez, 2006; Rueckl et al., 2015). Still several authors found, at the group level, that Chinese/Japanese character recognition induces slightly more right-lateralized VOTC activations than alphabetical reading (e.g. Bolger et al., 2005; Feng et al., 2020; Koyama et al., 2014; Qu et al., 2019; Szwed et al., 2014; Tan et al., 2001). We did not find such rightward bias, possibly because our participants were native Chinese speakers, while the rightward bias may be specific to logographic scripts acquired as a second language (for a review see Cao, 2016).

More generally, in the literature on bilingual reading, a tension between assimilation and accommodation has been proposed for the learning of the first (L1) and second (L2) languages (Perfetti et al., 2007). When L2 is similar to L1, the brain would utilize the same network of L1 for reading L2 (assimilation). This was shown in fMRI studies of English and Korean bilinguals where both languages are alphabetical (e.g. Kim et al., 2016). When L2 is very different from L1, the brain would use new areas for reading L2 (accommodation). This was suggested to occur for English L1 participants learning Chinese as L2, including in the right fusiform region (Y. Liu et al., 2007; Perfetti et al., 2007), middle occipital gyri (Kim et al., 2016), middle frontal gyrus (Y. Liu et al., 2007). Our results are partially in line with this theory, but that there is still much assimilation in English-Chinese readers, since many voxels show similar responses and word-similarity effects for both languages.

We see two reasons for the partially distinct cortical specialization in English-Chinese readers. The first hypothesis is that visual features of Chinese characters are different from those of the Roman alphabet used for English and French. While all writing systems share similar graphic principles and appeal to similar line intersections (Changizi et al., 2006; Changizi & Shimojo, 2005), they differ in the nature, number, and spatial arrangement of the shapes used, which may require a dedicated set of neural circuits. In support of this idea, we recently showed that, when recycling a convolutional neuronal network to recognize 1000 written French words, the network dedicates a few dozen units to letter shapes (Hannagan et al., 2021). In new simulations, we trained the same network to recognize either 500 English and 500 Chinese words, or 500 English and 500 French words, thus providing an elementary simulation of bilingual reading (Agrawal & Dehaene, unpublished simulations). The English-French network developed 54 word-selective units (±5 across replications), none of which were selective to a specific language. However, the English-Chinese developed 114 (± 5) word-selective units, out of which 26 (±5) were selective to Chinese characters and 14 (±4) to the alphabetic. Thus, for the purposes of an artificial network solely driven by the needs of invariant visual recognition, it is advantageous to dedicate some unique processing resources to the shapes of Chinese and alphabetic writing. It seems likely that the same constraints on invariant visual recognition would apply to the cortex.

Visual shape alone is unlikely to entirely explain ventral visual cortex specialization, however, because Arabic numerals share visual features with letters, and yet are processed in a specialized visual number area lateral to the OTS/IFS (Amalric & Dehaene, 2016; Daitch et al., 2016; Shum et al., 2013). Moreover, there is strong overlap of the VOTC regions activated during visual reading, and during reading through tactile or auditory channels in blind persons in previous studies at 3T (Reich et al., 2011; Striem-Amit et al., 2012). Thus, an alternative possibility is that the localization of the VWFA is heavily influenced by the prior connectivity of cortex to other areas (Bouhali et al., 2014; Dehaene-Lambertz et al., 2018; Hannagan et al., 2015; Saygin et al., 2016). Since Chinese is a less phonologically transparent language as compared to English and French, it may put heavier demands on the direct lexical route, thus relying on a slightly different set of connections and therefore, putatively, a different preferred site of origin in ventral visual cortex (Bouhali et al., 2019).

A limit of the present study is that we only studied two pairs of languages, thus leaving open many questions: Do partially different cortical patches take on reading in L1 and L2 only when the writing systems are very different, as is the case for Chinese characters vs Roman alphabet? Or would a similar split be seen for two different alphabets, for instance in Russian-English bilinguals? Does the direction of reading matter, as could be studied Arabic-French or Hebrew-English bilinguals? And, would VOTC also split its resources in bilingual readers mastering two languages with the same alphabet but very different letter statistics, such as French and Polish? Answering these questions would shed light on the causal factors underlying the cortical specialization for reading – but we should keep in mind the difficulty of recruiting a large group of such fluent bilingual volunteers for a 7T fMRI study.

### Sensitivity to the statistics of written language

Finally, the current study underlines the replicability of the word similarity effect (Vinckier et al., 2007), both with English-French alphabets, but also in Chinese (as recently reported by Guo et al., 2022). When written stimuli respect increasingly well the statistics of words in the known language, they induce increasingly greater activity in the VWFA (Binder et al., 2006). Many word specific clusters in English-Chinese participants even shared similar word-similarity profiles for English and Chinese. The present findings indicate that this property is not a general feature of the entire VOTC, as the heavily smoothed group-level images of Vinckier et al., (2007) may have suggested, but is a very local feature unique to word-specific patches and which is absent, for instance, in face-responsive patches that are not immediately adjacent to the word-responsive ones.

There are two complementary mechanisms which may give rise to this effect. First, in a bottom-up manner, during reading acquisition, the visual cortex may incorporate the statistics of the shapes (e.g. letters and letter groups) that make up words. Indeed, imaging data (Agrawal et al., 2019; Baker et al., 2007) and simulations (Hannagan et al., 2021) show how visual neural networks become progressively attuned to the specific shapes of letters and their combinations that are useful to recognize words in a specific script. As a result, already during the initial bottom-up wave of VOTC activation, the cortex would be increasingly responsive to strings of letters that increasingly mimic the statistics of real words. The second, not-necessarily exclusive possibility is a top-down mechanism: the visual inputs that are more similar to real words may be processed at a higher level, for instance evoking several candidates within the lexicon, and thereby elicit more top-down information from higher brain regions back to visual cortex (Hickok & Poeppel, 2007).

Owing to its poor dynamic resolution, fMRI is largely unable to separate those two possibilities. However, intracranial recordings suggest that both mechanisms may contribute, at different times following stimulus onset (Woolnough et al., 2021). Broadband gamma activity in left VOTC shows a larger initial peak (∼250 ms) when stimuli are more similar to real words (Woolnough et al., 2021), suggesting a bottom-up effect. However, this early effect is dominated by a difference between stimuli with frequent versus infrequent letters, suggesting that the bottom-up wave may primarily compile letter-based statistics. The effect of larger orthographic units, i.e. bigrams and quadrigrams, only appears later and in a top-down manner, seemingly triggered by a lexical effect first emerging in the anterior fusiform (Cohen et al., 2021; Woolnough et al., 2021).

Under the latter interpretation, while word-specific VOTC would be genuinely tuned to letters (contra Price & Devlin, 2011), most of the continuous, monotonic word-similarity effect seen here with fMRI would be due to top-down signals proportional to the strength of lexical activation. Such an interpretation is compatible with (1) simulations of purely bottom-up neural networks for reading, which only show a difference between frequent and infrequent letters, and not the full Vinckier (2007) word-similarity effect (Hannagan et al., 2021); (2) evidence for reversal of lexicality effects in VOTC when the task changes from passive viewing to active sentence reading, highlighting the influence of top-down processing on these activity profiles (Taylor et al., 2013; Woolnough et al., 2021); (3) evidence for top-down activation of VOTC during spoken language processing (Cohen et al., 2021; Dehaene et al., 2010, 2015; Yoncheva et al., 2010); and (4) the present evidence for greater VOTC activity for the non-dominant language in English-French bilinguals, which we interpret as a top-down effect due to greater attention and cognitive effort.

In conclusion, the present results illustrate the power of cortical plasticity and education in shaping the fine details of the human visual cortex. Not only do localized patches of cortex, at a millimeter scale, become highly attuned to the statistics of words and word-like stimuli in the learned script, but they may even become tuned to a single script in bilingual readers, at least when those scripts differ sufficiently at the visual level.

## Materials and Methods

### Participants

21 English-French bilinguals and 10 English-Chinese bilinguals were recruited from the Paris region and took part in this study.

The 21 English-French participants consisted of 3 sub-groups (7 participants each): a balanced early bilingual sub-group (age range 18-35, mean age=22.7, 4 females), where participants grew up in bilingual environments and were native speakers/readers of both English and French, having acquired both languages before the age of 10; an English-dominant sub-group (age range 21-32, mean age=25.4, 2 females); and a French-dominant sub-group (age range 20-26, mean age=22.1, 4 females). In the latter two sub-groups, participants acquired one of the languages as their native language, and later learned the other language at school, such that they became fluent readers (see below).

Given our testing site (NeuroSpin, near Paris), and the fact that those experiments took place during the COVID epidemic, we could not compose similar sub-groups for English-Chinese bilinguals (indeed, participants who learn Chinese as a second language at school and yet become fluent readers are quite rare). Instead, the 10 English-Chinese participants (age range 20-31, mean age 25.7, 7 females) were all native speakers/readers of Simplified Chinese, and later became fluent in English.

All participants were assigned to groups according to their self-reports and the Language Experience and Proficiency Questionnaire (Marian et al., 2007) that they filled in during recruitment. Participants also completed an online language test for English and French (https://dialangweb.lancaster.ac.uk/getals, tests on listening and structure, the English-Chinese participants only completed the English test). All achieved at least B2 level. A few participants were also familiar with a third/fourth language, although the proficiency was always at a much lower level comparing to the 3 languages tested in the current study.

All participants were right-handed, had normal or corrected-to-normal vision, and no histories of neurological/psychiatric disorders or reading/learning difficulties. Participants provided written consents for the fMRI study, and received monetary compensation. The study was approved by the local ethical committee in the NeuroSpin center (CPP 100032 and 100055) and the study was carried out in accordance with the declaration of Helsinki.

### Stimuli

#### Functional localizers

The stimuli for the English-French functional localizer (here-on abbreviated as “localizer”, **Figure 1A**) were gray-scale images from 8 different object categories (20 exemplars per category). They were adapted from stimulus databases and previous experiments, including faces (neutral, 10 males) (Lundqvist et al., 1998), bodies without heads (neutral standing still, 10 males) (de Gelder & Van den Stock, 2011), English words (Keuleers et al., 2012), French words (http://www.lexique.org/) (New et al., 2004), numbers (famous mathematical constants, 6-7 digits, including the decimal point and +, - signs), false fonts (6 letters, no letter was repeated on the same position) (Vinckier et al., 2007), houses, tools (half in a position graspable by the left hand). Both English and French words were 6-letter strings, with word frequency (computed from film subtitles) ranging from 7 to 761 per million. Words and numbers were rendered in the font Consolas. 9 words and 12 numbers were adapted from the category localizer in (Amalric & Dehaene, 2016). Houses and tools were adapted from the category localizer in (Zhan et al., 2018). All stimuli had a width or height less than 227 pixels (the longer axis less than 241 pixels) and were embedded within a gray circle (RGB: 157, 157, 157; diameter=310 pixels, 3.2°) presented at the center of a black screen. Stimuli were controlled across conditions for mean luminance and mean root-mean-square contrast within the circle. In an additional checkerboard condition, two alternating checkerboards of opposite pixel intensities were presented at the same temporal pace as the other stimuli categories, filling in the whole gray circle. A black star served as the target, the participants were required to press a button with the right thumb on a cylinder button box as soon as they detected it.

Most of the conditions in the English-Chinese localizer were identical to the English-French localizer. Only the French words were replaced by Chinese two-character real words (rendered in the font Source Han Mono which was very similar to Consolas, word frequency matched to the English words)(Cai & Brysbaert, 2010), and the checkerboards were replaced by the single strokes decomposed from the Chinese real words.

#### Main fMRI runs

### English-French stimuli

In order to build the English-French materials, we first computed the (log) frequencies of letters, bigrams and quadrigrams in English and French, using lexical frequency information from the SUBTLEX field from the British Lexicon Project database (Keuleers et al., 2012), http://crr.ugent.be/programs- data/lexicon-projects), and the FREQFILMS field from the French Lexique database (New et al., 2004), version 3.82, https://chrplr.github.io/openlexicon/datasets-info/Lexique382/README-Lexique.html), Secondly, we generated all possible 6-letter strings using the 26 letters of the alphabet (ignoring French accented letters). We then excluded all strings which did not include a single quadrigram existing in French or in English, thus reducing the corpus by a factor of ∼10. In order to avoid spurious lexical effects, we also removed strings in which 5-letters words were embedded. For each remaining 6-letter strings, we obtained the average frequencies of its letters, bigrams and quadrigrams, in English and French.

From this set of 6 letters strings, we created 14 categories of stimuli, each comprising 180 items (**Figure 1B**). Four categories, devoted to the study of letter frequency, resulted from the crossing of French statistics (strings with high-vs. low-frequency letters) x English statistics (strings with high-vs. low-frequency letters). Four similarly crossed categories were devoted to the study of bigram frequency, and 4 others to the study of quadrigram frequency. All stimuli so far were nonwords. The two final categories, devoted to the study of lexicality, consisted in real French and English words.

For the 12 sublexical components conditions, we used only nonwords, and selected sets of stimuli so that frequency should not be correlated across types of sublexical components (**Figure 1C**). For instance, we wanted to make sure that effects of quadrigram frequency could not result from effects of bigram frequency, although the frequency of quadrigrams and bigrams tend to be correlated in random letter strings. The selection of stimuli was optimized based on the average frequencies of its component letters, bigrams, and quadrigrams (**Figure 1C**).

For the two lexical conditions, we selected French words which were not English words, and conversely. Those words were matched in all respects (frequency of letters, bigrams, and quadrigrams) with stimuli from the quadrigram categories: English words with items with high frequency of quadrigrams in English and low frequency in French, and conversely for French words. This amounted to comparing real French words to French-looking pseudowords, and the same for English.

The code for generating the stimuli can be found at: https://github.com/chrplr/compute-pseudowords.

All stimuli were rendered during the experiment using the monospaced font Consolas at 66 points.

### English-Chinese stimuli

#### English stimuli

The 7 English stimuli conditions were taken from the English-French experiment, i.e. 6 word component conditions with low or high frequencies in both languages (L-, L+, B-, B+, Q-, Q+) plus English real words.

#### Chinese stimuli

In the Chinese writing system, words are formed by one or more characters (in a specific order; changing the order often changes the meaning of the word). The characters themselves are composed of one or several radicals (graphical components) in different orthographic compositions (e.g. left-right, up-down). Some radicals are themselves compounds of radicals that can be further decomposed. Finally, all radicals can themselves be decomposed into several elementary writing strokes. Given this organization, by analogy to the English and French stimuli, the first author (M.Z.), a native speaker of Chinese, generated a hierarchy of 7 Chinese-stimuli conditions that all spanned the space of two Chinese characters but were increasingly similar to real Chinese words. Each condition contained 180 stimuli. Note however, that the 7 Chinese conditions were not directly parallel to the 7 English conditions.

The top 3 conditions in this hierarchy (WH, WL, CP, see **Figure 5**) all contained real Chinese Characters with matched frequencies. We constructed 180 high-frequency two-character real Chinese words (condition WH; log 10 frequency: 1.5 to 2, mean=1.744) from 196 unique characters (all high frequency, log 10 frequency ranging from 2.5 to 3.5). The low-frequency real words (condition WL; log 10 frequency: -0.5 to 0, mean=-0.217) were then composed of 158 of the 196 abovementioned unique characters. Finally, the character pair condition (CP) were generated from the 185 of the 196 unique characters in condition WH, excluding combinations that (1) were real words, (2) would become a real word if the two characters were flipped, (3) had a pronunciation similar to a real word, (4) were often contiguous characters in daily-life texts (e.g. two connecting characters from two adjacent words).

Two other conditions contain Chinese radicals in orthographically possible (condition RP) and impossible (condition RI) positions, respectively. Condition RP was generated by decomposing all 360 real characters from condition CP into radicals, and reassembling them in orthographically possible positions, with a slight change in visual form when appropriate (in Chinese calligraphy and typography, the form of a given radical can subtly change when used in different sizes and positions; e.g. the characters 石磊蠹 share the same radical 石 in different positions). For Chinese readers, the resulting RP stimuli were perceived as extremely low-frequency real Chinese characters that did not remind them of common real words. We checked that RP stimuli were absent from GB2312, GBK, or GB18030 Chinese encoding standards (∼21,887 ideographs); the Source Han mono font’s CJK glyph database (48,966 ideographs); and the character decomposition series in zh.wiktionary.org; but could exist in Unicode’s CJK Unified Ideographs Extensions (92,856 ideographs), which includes non-Chinese (Japanese or Korean) characters.

Condition RI contained exactly the same number and types of radicals as in condition RP, although shifted into orthographically impossible positions. They were made with the following guidelines. Whenever possible, radicals were placed at positions where they cannot appear in the Chinese script -- or only appear with very low probability i.e. very low character frequency. For “compound radicals”, which may still be further decomposed into sub-components e.g. two radicals, the sub-components themselves were switched (but such a reconstructed compound radical was occasionally kept at its original location). Finally, the characters we chose contained some high-frequency radicals which can appear at any possible positions (e.g. the radical 口, which appears in 吅吕品㗊哀回器… covering all orthographically possible positions; several other radicals with similar properties include:日又由火木且求). In most of these cases, locations for these radicals were left blank; in a few cases we chose a specific visual form of the radical (in the sense of calligraphy and typography) which could not appear at the chosen location.

Finally, the lowest-level conditions (S and SG) comprised only elementary strokes, directly decomposed from the high-frequency real-word characters (WH). For the first condition (S: strokes), all 360 characters of condition WH were broken down into Chinese GB13000.1-standard strokes in their natural forms, conforming to the rules of Chinese handwriting (recovering the crossings and corners formed by overlapping strokes, no artificial stroke breaks, no stroke rotation). The strokes were rearranged so that they spanned the same overall two-character space, without intersecting, and without forming two blocks. The second condition of stroke groups (SG) contained the same strokes, except that the strokes formed two character-like blocks, yet without forming any real Chinese radical or character (at least in their exact visual forms in the Source Han mono font). The number of crossings, connections and overlapping corners was kept very similar to the high-frequency real-word condition (WH).

#### Rendering

The English and Chinese stimuli were both rendered in the monospaced font Source Han Mono (https://github.com/adobe-fonts/source-han-mono), whose strokes are visually similar to the alphabetical font Consolas. English fonts were rendered at 56 points and regular weight (corresponding to the size of 66 points in Consolas), Chinese fonts were rendered at 71.27 points with normal weight, so that the stroke thickness and spatial areas covered by the two characters were roughly matched to the 6-letter English strings.

The Chinese stimuli were decomposed and recombined in vector forms in Adobe Illustrator CS 6.0. All stimuli were converted into images to make sure the font rendering remained exactly the same for all participants. We controlled the mean pixel values within the English and Chinese conditions. For the first 3 English conditions, we slightly changed the black pixel values (RGB 0, 0, 0 to 2, 2, 2) for condition L-, and made some of the anti-aliased edge-pixels on the letters slightly darker for condition L+ and B-. This resulted in matched mean pixel values across all English conditions, without noticeable change in the visual percept. For the first 4 Chinese conditions, we made the strokes 1 pixel thinner for the same reason.

### Data acquisition

The brain images were acquired using a 7-Tesla Magnetom scanner (Siemens, Erlangen, Germany) with a 1Tx/32Rx head coil (Nova medical, Wilmington, USA) at the NeuroSpin center of the French Alternative Energies and Atomic Energy Commission (CEA). Dielectric pads were used for 30/31 participants (not used for SE01 due to insufficient space within the head coil). To minimize head movements, a tape (padded with tissue papers for comfort) was fixed on the forehead of each participant and the head coil, to give tactile feedback to the participants whenever they attempted to move their heads (Krause et al., 2019). To minimize light reflections inside the head coil, a piece of black paper was inserted to cover the inner surface of the transmitter coil element. Stimuli were presented via a BOLDscreen 32 LCD screen (Cambridge Research Systems, Rochester, UK, 69.84 x 39.29 cm, resolution=1920 x 1080 pixels, refresh rate=120 Hz, viewing distance ∼200 cm), at the head-end of the scanner bore. Participants viewed the screen through a mirror attached to the head coil. The whole scanning session lasted about 90 minutes.

Within each experiment, one localizer run (9 min 12 s, 276 volumes) and 3 main fMRI runs (13 min 18 s per run, 399 volumes) were acquired (due to scanner technical issues, only two main fMRI runs were acquired for participant SB01). Functional data were acquired with a 2D gradient-echo EPI sequence (TR = 2000 ms, TE = 21 ms, voxel size = 1.2 mm isotropic, multiband acceleration factor=2; encoding direction: anterior to posterior, iPAT=3, flip angle = 75, partial Fourier=6/8, bandwidth=1488 Hz/Px, echo spacing=0.78 ms, number of slices=70, no gap, reference scan mode: GRE, MB LeakBlock kernel: off, fat suppression enabled). To correct for EPI distortion, a 5-volume functional run with exactly the same parameters apart from an opposite phase encoding direction (posterior to anterior) was acquired immediately before each task run. Participants were instructed not to move between these two runs. Manual interactive shimming of the B0 field was performed for all participants. The system voltage was set to 250 V for all sessions, and the fat suppression was decreased per run to ensure the specific absorption rate (SAR) for all functional runs did not surpass 62%. To minimize artifacts and increase signal-to-noise (SNR) ratio around the VOTC, for 28/31 participants, the functional data acquisition slab was placed in a way that excluded the eyes and the ear canal signal dropout region, so that the VOTC, especially the anterior occipital-temporal sulcus (OTS) above the ear canal, was covered as much as possible (see **Figure S1**). The ear canal signal dropout only affected the anterior VOTC data of the first 3 participants (SB01, SB02, SE01), but did not affect the more posterior location of the classical VWFA.

High-resolution MP2RAGE anatomical images were obtained after 2 or 3 functional runs (resolution=0.65 mm isotropic, TR=5000 ms, TE=2.51 ms, TI1/TI2=900/2750 ms, flip angles=5/3, iPAT=2, bandwidth=250 Hz/Px, echo spacing=7 ms).

### Experimental design

#### Functional tasks

Both the localizer and the main fMRI runs used a mini-block design, where multiple stimuli were presented rapidly in each block. A green fixation dot (RGB: 112, 219, 96, dot diameter=8 pixels) was always present at the center of the screen. Participants were required to detect a rare target which, in some of the blocks, randomly replaced a stimulus after the 5^th^ within a block.

For the localizer, each block contained 20 stimuli and lasted for 6 s, followed by a jittered fixation period of 4, 6, or 8 s (mean=6 s). Within each block, the stimuli were presented for 100 ms, followed by a fixation period of 200 ms. Each of the 9 experimental conditions was repeated for 5 times, 2 of which containing catch trial targets, except the checkerboard condition which didn’t contain catch trials. The target of the catch trials was a star. Additional fixation periods of 6 s were added to the start and the end of each run.

For both the English-French and English-Chinese main fMRI runs, each block contained 12 stimuli and lasted for 4.2 s, followed by a jittered fixation period of 3.8, 5.8, or 7.8 s (mean=5.8 s). Within each block, the stimuli were presented for 150 ms, followed by a fixation period of 200 ms. Each of the 14 experimental conditions were repeated for 15 times, randomized and counterbalanced within 3 runs (5 repetitions per run, 2 of which containing a catch trial target). Across languages, the target was always 6 hashtags (######). The stimuli within each condition were only presented once, and the presentation order within each block was fixed, to prevent the same letter appearing in the same position in consecutive stimuli for the English-French experiment. Each run contained 70 blocks of stimuli, and an additional 7 blocks of fixation-only blocks (jittered 8, 10, 12 s, mean=9.71 s). Fixation periods of 16 s and 14 s were added to the start and the end of each run. The background color of the screen was gray (RGB: 128, 128, 128).

#### One-minute word reading test

Immediately after the scanning session, the participants performed a one-minute word reading test for each language (English-French participants: English and French lists; English-Chinese participants: English, Chinese, French lists). Each list contained 160 high-frequency words. The English and French words had 5-8 letters; the Chinese words had 2 characters. The words did not overlap with the ones in the main fMRI runs. Participants were instructed to read the words aloud, as fast as possible but also as clearly as possible. The order of languages was randomized for each participant. The number of words read within one minute was recorded for each language.

A language dominance score was computed for each participant, based on their number of words read for the corresponding two languages tested in the main fMRI runs (abbreviated here as L and L’): [number of L words - number of L’ words]/[total number of L and L’ words]. The resulting scores corresponded well with the participants’ self-reported language history and dominance. The scores of the English-French participants lied on a continuum, without clear boundaries between the early bilingual, English-dominant and French-dominant sub-groups (see **Figure S4**). For English-Chinese participants, however, the dominance was always Chinese > English > French (Chinese-dominant).

### Data analysis

The data were analysed in BrainVoyager v21.45 (Brain Innovation, Netherlands), Matlab R2018b, and the Matlab package NeuroElf v1.0 (https://neuroelf.net/).

#### Data preprocessing

The functional data underwent top-up distortion correction (COPE plugin in BrainVoyager), where the in-plane voxel displacement map between the first volumes of both the actual task run and its corresponding distortion correction run was computed, and applied to the task run. The distortion-corrected data was then corrected for slice scan time (sinc interpolation, slice scanning order read from the slice time table in the DICOM headers), 3D rigid motion correction (trilinear for estimation, sinc for applying the correction, aligned to the first volume within each run), high-pass temporal filtering (GLM with Fourier basis set, number of cycles=2). No spatial smoothing was applied to the data at this stage.

The MP2RAGE anatomical data consisted of four image types: inversion 1, inversion 2, quantitative T1, uniform T1w. To have a similar appearance to the conventional MPRAGE anatomical data, the uniform image was divided by the T1 image (this step is optional), and the background noise was masked out by the inversion 2 image. The resulting anatomical image was resampled to 0.6 mm isotropic (framing cube dimension: 384×384×384), and transformed into Talairach space (TAL). For data visualization in **Figure S2 and S5**, the white matter-grey matter boundary was segmented in TAL space, and reconstructed as surface meshes.

For fMRI across-run co-registration, we used a manual procedure and achieved better co-registration qualities than the automatic procedure: the localizer run was co-registered to the anatomical data, then all the other functional runs were manually co-registered to the localizer run. For quality checks, the across-run co-registration quality was inspected visually with animations looping through the first volumes across runs in TAL space. By co-registering with the same type of image modality (T2*), any mismatch between two runs (even in cases below 0.1 mm or 0.1 degree) became extremely easy to spot and correct. After the quality checks, all functional images were then transformed into TAL space and were kept at 1.2 mm isotropic resolution.

#### Statistical analysis

All statistical tests in this study were two-tailed, including whole-brain contrasts.

### Whole-brain general linear model (GLM) analysis

In individual analysis, for both the localizer and the main fMRI runs, the blocks of conditions plus button presses for target trials were defined as predictors and were convolved with a canonical two-gamma hemodynamic response function (HRF); the 6 head-motion parameters were z-scored and defined as confound predictors. The target stimuli (stars and “######”) were not modelled, since they were collinear with the button presses. The time-course data underwent %-transformation before performing the fixed-effect GLM, and the serial correlations in the data were corrected with an AR(2) model.

For the localizer, the group-level GLM was performed with the participants as the random effect, with both unsmoothed data and data smoothed with a Gaussian filter of 6 mm full-width-at-half-maximum (FWHM). Whole-brain contrasts were initially thresholded at p<0.001, then underwent Monte-Carlo simulation for multiple-comparisons correction (Forman et al., 1995) using the Cluster-level statistical threshold estimator plugin in BrainVoyager software: 5000 samples of null data with the same map smoothness as real data were generated, and the contiguous cluster size that occurred in null data with an overall frequency lower than 0.05 (alpha<0.05) was set as the cluster size threshold. For descriptive and visualization purposes (**Figure 2B**), the event-related average time course per condition for the group-level cluster (and individual clusters/ROIs) were computed, subtracted and divided by the baseline (activity of -2 to 0 TRs before each block).

Functional data smoothing was only applied to the group-level whole-brain analysis; no data smoothing was applied to subsequent individual-level analyses, either at the whole-brain level or at the individual ROI level.

### Individual ROI definition

For each individual participant, the word- and face-specific clusters in the localizer were defined as ROIs (bilingual word contrast: average of English and French words > faces, bodies, houses, tools; face contrast: faces > bodies, houses, tools, average of English and French words; p<0.001 uncorrected, trilinear interpolation, cluster size > 4). Two additional single-language-specific contrasts were computed (e.g. English > faces, bodies, houses, tools) for both groups to examine if the bilingual word-specific contrast included most of the language-specific voxels, which was the case for the English-French group but not for the English-Chinese group. Thus to maximize sensitivity, for English-Chinese participants, the language-specific (Chinese-specific and English-specific) ROIs were defined by the main fMRI runs, which had much more block repetitions per condition than the English and Chinese word conditions in the localizer data (15 compared to 5 repetitions). We used the positive and negative clusters from the contrast of average activity evoked by high and low-frequency Chinese words versus activity evoked by English words, p<0.001 uncorrected, trilinear interpolation, cluster size > 4. A similar analysis was attempted for French versus English words in English-French participants, but it yielded few clusters that were not reproducible across the localizer and main fMRI runs (see main text). To avoid the possibility that we missed voxels sensitive to English or French only, in a supplementary analysis we computed contrasts for single languages (e.g. English words only > faces, bodies, houses, tools), sorted voxels according to whether they overlap with the bilingual word ROIs or not, and examined the single-condition activity profiles across languages within each kind of voxels. The activity profiles were again inconsistent between the localizer and the main fMRI runs for English-French participants.

Throughout the study, the conditions used to examine ROI properties were independent from the conditions used to define the ROIs (conditions from different types of runs i.e. the localizer versus the main fMRI runs, or different conditions within the same runs).

For each ROI, the broad anatomical region it belonged to (ventral occipito-temporal, lateral temporal, lateral frontal, IPS, medial frontal, others) and the exact anatomical location were manually labelled, according to (Rossion et al., 2018) and (Duvernoy, 1999). For VOTC ROIs, the sub-regions they belonged to (EVC, fusiform) were further labelled; and the fusiform sub-region was even further split into IOS-OTS, mid-fusiform sulcus (mFS), collateral sulcus (CoS), middle temporal/occipital sulci/gyri (MTS/MTG/MOS/MOG) according to individual anatomy. Clusters outside the brain, in the cerebral spinal fluid or in the white matter were excluded as outliers. We also observed activation clusters having a shape of blood vessels (clearly seen from their 3D surface reconstructions). They overlapped or were adjacent to blood vessels, and yet some showed condition-specific activity with normal BOLD HRF shapes. The reason of their occurrence in 7T fMRI data is not yet clear, and could potentially be caused by the shift of B0 around venous blood vessels, (Winawer et al., 2010), a hypothesis which needs further examination in future studies. In the current study, we excluded most of these vessel-shaped clusters, and included only the ones with much bigger cluster sizes than the vessels and having normal HRF shapes.

All ROIs after outlier exclusion were visualized as 3D surface meshes, see **Figures S2 and S5**. For each ROI, the averaged x, y, z TAL coordinates across voxels were extracted for figure plotting and subsequent analyses. The single-condition beta values (% signal change versus fixation) of the localizer and main-fMRI-run GLMs were also extracted per voxel, and averaged per ROI.

For both the English-French group and the English-Chinese group, to examine the laterality of word- and face-specific clusters in the VOTC, the number of clusters and voxels were counted for each individual participant, and submitted to a paired t test.

### Word similarity effects in ROIs

For English-French participants, we fitted the averaged beta values for the 14 main conditions with the linear predictor of [1 2 2 3 4 5 5 6 7 8 8 9 10 10]/10 (normalized between 0 and 1 by dividing with 10), which assumes higher activity for higher frequencies within each word component (low English-low French frequency vs. high English-high French frequency conditions within letters, bigrams, quadrigrams), and higher activity for components more similar to words (letters<bigrams<quadrigrams<words), but does not assume a difference in activity between languages (LE=LF, BE=BF, QE=QF, WE=WF). The significance of the linear fit was corrected for multiple comparisons with FDR correction (BH procedure, https://www.mathworks.com/matlabcentral/fileexchange/27418-fdr_bh) across all participants and all ROIs, or for ROIs within specific brain regions. The slope of the fit per ROI was used as a measure of the word similarity effect for further analysis.

For English-Chinese participants, the linear predictor [1 2 3 4 5] (normalized between 0 and 1) was separately fitted to the first 5 English and Chinese conditions (avoiding the last two Chinese conditions, because those were used to define language-specific ROIs). No ROI survived FDR correction, likely due to the much fewer conditions for fitting here compared to the English-French experiment (5 vs 14), instead of any quantitative differences in the data. We validated this possibility by fitting the 5 corresponding English conditions in the English-French participant data, resulting in a similar level of significance (0/773 ROI survived FDR correction, comparing to 499/773 ROIs when fitting all 14 conditions). Thus we did not report FDR correction here for the English-Chinese participants. Also, because there were no explicit correspondence of the word component levels between English and Chinese conditions, we did not directly compare the English and Chinese slopes in the same participants.

### Word and other functional gradients across VOTC word-specific ROIs

We examined the gradual posterior-to-anterior changes of multiple functions across VOTC ROIs (such changes across cortical spaces were termed “gradients” here). Within each individual participants across bilateral VOTC ROIs, one specific functional value per ROI was fit to the TAL Y coordinates of the ROI, resulting in one regression coefficient per participant. A significant fit indicates a posterior-to-anterior change of that function. When examining the laterality effects of the gradients, the left- and right-hemisphere ROIs were fitted separately. The regression coefficients across participants then underwent a t test (one-sample t tests against 0 to examine the significance of the gradients, and paired t tests between left and right hemispheres). For statistical tests involving the right-hemisphere ROIs, only participants with at least 3 right-hemisphere ROIs were included.

The examined functional gradients included: (1) word similarity slopes (separately for each hemisphere); (2) activity evoked by English words, French words, false fonts and numbers (across bilateral ROIs); (3) the word selectivity index (separately for each hemisphere), computed as [word activity - other activity]/[word activity + other activity], where word activity = averaged activity of English and French words; and other activity = averaged activity of faces, bodies, houses, tools. We followed a padding procedure to ensure that the word selectivity index ranged from -1 to 1 (Simmons et al., 2007): if the activity in the condition with the smallest amplitude was negative, all conditions were padded with the absolute activity value of this negative condition, so that the activity for all conditions was zero or positive.

(1) was computed with the main fMRI runs data, for both the English-French and English-Chinese groups. A Kruskal-Wallis test was further performed to compare the English word gradients in word-specific ROIs between the two groups of participants. (2) and (3) served to further characterize word selectivity in VOTC, especially for ROIs in the anterior fusiform. They were computed with the localizer data, and only for the English-French participants.

### Analyses of main-fMRI-run conditions in word-specific ROIs

With GLMs and contrasts performed on the averaged time courses of each word-specific ROI, we examined the main-fMRI-run conditions in more detail, aiming to see whether the potential lexicality effect, frequency effect, and language differences were associated to specific anatomical locations. See **Figure 1 and 5** for condition abbreviations.

For English-French participants, we examined the lexicality effect using the contrast [WE, WF > QE, QF]. Within each word component, we also examined the component frequency effect, contrasting between conditions having high frequency versus conditions having low frequency in both languages (L+ > L-, B+ > B-, Q+ > Q-). We further compared the difference of these frequency and lexicality effects between adjacent component levels ([B+ > B-] > [L+ > L-], [Q+ > Q-] > [B+ > B-], [WE, WF > QE, QF] > 2[Q+ > Q-]) to see if there were increased sensitivity for specific word components. To examine the English-French language differences, we contrasted the conditions having high frequency in one language within each word component (LF > LE, BF > BE, QF > QE). We also submitted the 2×2 data from each word component to a standard analysis of main effects and interaction. Taking the letter component for example, the main effect of French was examined by the contrast [LF, L+ > LE, L-]; the main effect of English was examined by [LE, L+ > LF, L-]; and the interaction of French and English was examined by [L+ > LF] > [LE > L-].

In English-Chinese participants, for the English conditions, we performed the same frequency effect contrasts (L+ > L-, B+ > B-, Q+ > Q-), and the contrast WE > Q+ for the lexicality effect. For the Chinese conditions, we contrasted adjacent conditions, including contrasts between non-character conditions (SG > S, RI > SG, RP > RI), real characters > non-characters (CP > RP), and contrasts involving real words (WL > CP, WH > WL).

### Effect of individual language dominance

Separately within the English-French and English-Chinese groups, we examined whether the individual language dominance score (see the section “One-minute word reading test”) were correlated with brain activity differences between languages. Take English and French for example, the language dominance score was calculated as [numbers of English words - numbers of French words]/[total number of English and French words].

In the brain of individual participants, we first merged ROIs within each of the 3 broad regions (Corresponding to regions in **Figure 3**, and details described in the section of “Individual ROI definition”). Furthermore, we merged VOTC ROIs into sub-regions including EVC, fusiform, and even small sub-regions in the fusiform including the IOS-OTS sub-region, and the mFS sub-region. We then averaged the single-condition beta values within each region or sub-region, and computed the activity difference between English and French conditions (English-French group: localizer: WE - WF; main fMRI runs: LE - LF, BE - BF, QE - QF, WE - WF; English-Chinese group: WE - WC in both the localizer and the main fMRI runs, false fonts - Chinese strokes in the localizer). Then these activity differences for each region were correlated to the language dominance scores across participants.

### Non-parametric ICA analysis decomposing sub-voxel components in word-specific ROIs

We used the code for non-parametric ICA analysis from (S. Norman-Haignere et al., 2015) (https://github.com/snormanhaignere/nonparametric-ica), which maximizes the non Gaussianity and finds independent components in the data. The English-French and English-Chinese data were analyzed separately. The input data were voxel beta values for the 14 conditions in the main fMRI runs from all bilingual word-specific ROIs (the ones shown in **Figure 3** and **Figure 7**, defined by the localizer). For English-Chinese participants, removing the overlapping voxels showing Chinese or English language specificity did not change the results. We searched for 3 components, to allow us to separate the effects of the baseline, the word similarity effect, and putative language differences. Searching for more components resulted in the same 3 stable components, plus some additional unstable components which were much harder to interpret. We only report the 3 stable components here.

This analysis resulted in 3 component profiles per voxel, each with a corresponding weight. The observed data could be reconstructed by multiplying the component profile and the weights. To deduce the localizer activity profile corresponding to each component, we divided the observed localizer data by the main-fMRI-run component weights (**Figure 8**).

## Acknowledgments

We thank Alexandre Vignaud for the help in setting up the 7T scanning sequences; the nurses and MR-technicians at NeuroSpin for the help in participant recruitment and scanning; Aakash Agrawal for running simulations on bilingual neural networks. MZ is supported by Fondation Bettencourt Schueller, and Programme Investissements d’Avenir IHU FOReSIGHT (ANR-18-IAHU-01). LC is funded by the Programme Investissements d’avenir (ANR-10-IAHU-06) to the Paris Brain Institute. LC and SD are supported by the “TOPLEX” ANR program (ANR-20-CE37-0002).

**Conceptualization**, M.Z., S.D., and L.C.; **Funding Acquisition**, S.D. and L.C.; **Data Curation**, M.Z.; **Methodology**, C.P. and M.Z.; **Formal Analysis**, M.Z.; **Investigation**, M.Z.; **Project Administration**, S.D.; **Supervision**, S.D.; **Resources**, S.D.; **Software**, C.P. and M.Z.; **Validation**, M.Z.; **Visualization**, M.Z.; **Writing – Original draft**, M.Z., S.D., and L.C.; **Writing – Review & Editing**, M.Z., C.P., S.D., and L.C.

The authors declare that they have no competing interests. All data needed to evaluate the conclusions in the paper are present in the paper and/or the Supplementary Materials. The experimental stimuli and ROI data can be found at https://osf.io/96syx/?view_only=88b55034027042fbb4f118f398fc5706. The code for generating the English-French stimuli can be found at: https://github.com/chrplr/compute-pseudowords.

## Supplementary Materials

### Supplementary Text

In the main text for the English-French participants, we analyzed the activity in word-specific clusters irrespective of language (denoted as “bilingual clusters” below), defined by greater activity to the average of two languages (English and French) relative to other categories (faces, bodies, houses, tools) in the localizer runs (p<0.001 uncorrected, cluster size > 4). This was motivated by the finding that there were no consistent language-specific clusters when directly contrasting English and French words, neither in the localizer nor in the main fMRI runs, and most of the language-specific voxels were included by the bilingual voxels (whole brain, English: 91.90%, French: 86.14%), especially in the VOTC (English: 95.75%, French: 94.82%). However, it may seem strange to pull together the two languages, and only then check whether they prefer one language over the other, and may miss language-specific clusters that didn’t survive thresholding under direct comparison.

In this supplementary analysis, we show that this choice of examining word-specific clusters without language specificity (bilingual clusters) in the English-French participants was inconsequential, because it did not lead to missing voxels that would be putatively strongly selective to one language. Here we performed two separate contrasts with each single language (English words > faces, bodies, houses, tools; French words > faces, bodies, houses, tools, p<0.001 uncorrected, cluster size > 4), separated all voxels into four categories, according to whether they overlapped with the bilingual clusters, and examined the averaged activity within each voxel category. If a language-specificity was present, it would be consistently observed in the language-specific conditions for word components (LE vs LF, BE vs BF, QE vs QF) and for the real words (WE vs WF). We performed this analysis in a VOTC mask (**Figure S3A**), and also in the rest of the voxels outside the VOTC mask (**Figure S3B**). To minimize the possibility that potential language dominance difference between participants may obscure the language-specificity activity differences, we performed the analysis separately for the three sub-groups. As a comparison, we also performed the same analysis for the English-Chinese participants.

For English-French participants, both the group-averaged and single-subject results showed that there were no consistent language-specificity for any of the three sub-groups in the single-language-specific voxels (third and fourth columns). I.e. the direction of language specificity for word-components (LE vs LF, BE vs BF, QE vs QF) did not correspond to that of the real word conditions, and the real word conditions in the main fMRI runs were not even consistent with the ones from the localizer. Again this supports our finding in the main text that the English-French participants use the same set of brain areas to process the written scripts of both English and French.

On the other hand, the language specificity of English-Chinese participants was very consistent between the localizer and the main fMRI runs, no matter whether or not they were overlapping with the bilingual clusters. The single-language-specific non-overlapping voxels showed language specificity (higher activity amplitude), consistent with the finding that the bilingual word-specific voxels did not include all language-specific voxels. The slopes in the VOTC mask across conditions were also consistent with what we reported in the main text: In the bilingual clusters, a slope was both present for English and Chinese; in the Chinese-selective clusters though, a slope was only present for Chinese but not for English.

### Supplementary Figures

**Figure S1.**
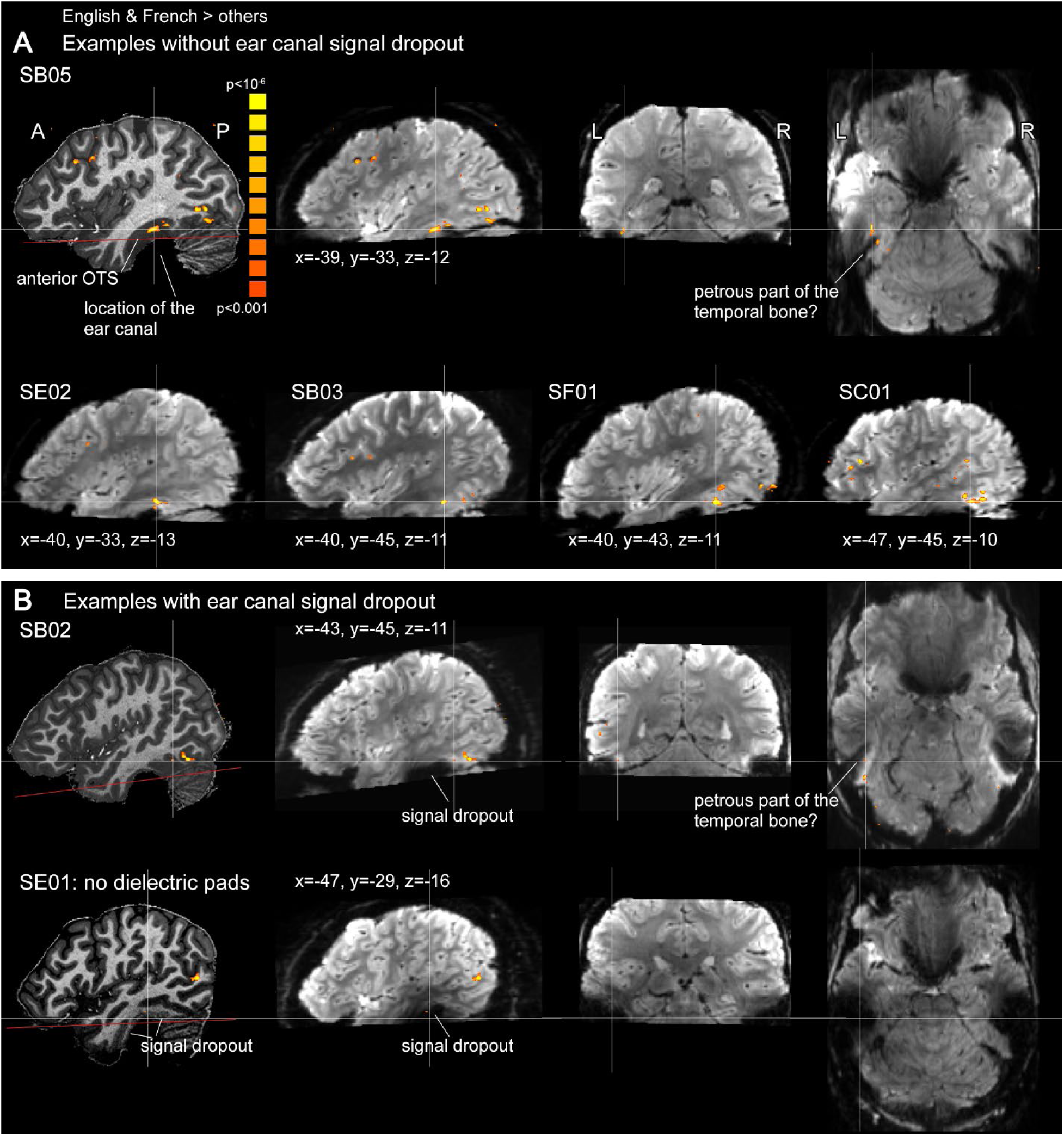
Functional data coverage and optimization for the ear canal signal dropout. The coverage for the majority of the datasets (28/31 across the two experiments) avoided the ear canal signal dropout. Panel **A** shows three orthogonal views from one participant (SB05), and sagittal views from one representative participant per group (SE02, SB03, SF01, SC01). Anterior VOTC word-specific activation clusters could be observed in these participants. A minority of datasets (3/31) were affected by signal dropout (SB01, SB02, SE01). Panel **B** shows the latter two participants. Note that even in this case, the clusters in the posterior VOTC (locations around the classical “VWFA”) were always present (TAL Y < -50, see **Figure S2**). The signal dropout was either caused by the inclusion of the ear canal into the data acquisition slab (SB01, SB02), or by not being able to use the dielectric pads in the tight space of the head coil (SE01). The subsequent reduction in SNR in lateral temporal cortices can be observed in both the anatomical and functional images. In both A and B, red lines overlaid on anatomical images indicate the bottom edge of the data acquisition slabs, and were parallel to the in-plane phase encoding direction.

**Figure S2.**
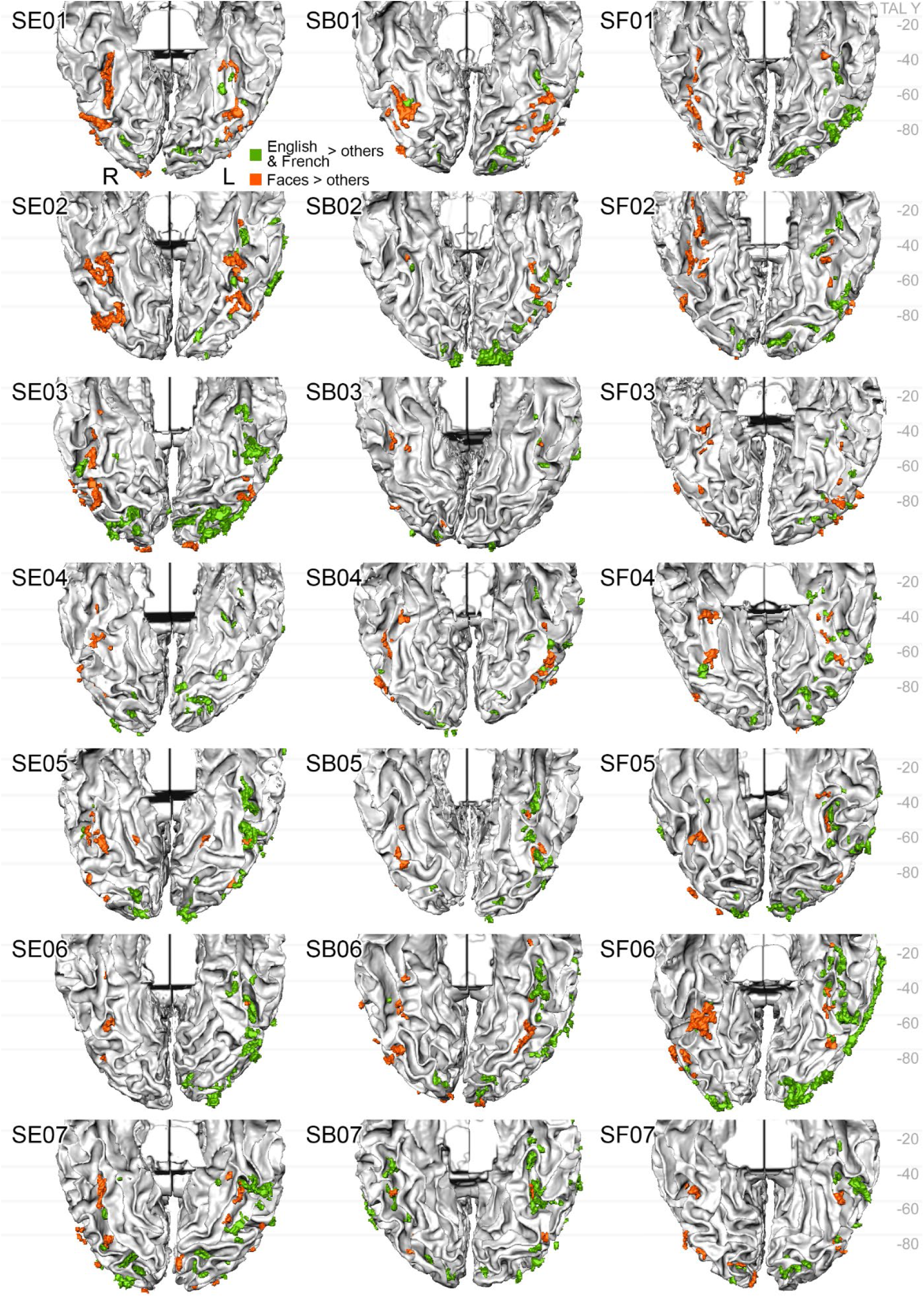
Word- and face-specific clusters in VOTC in each individual English-French participant. (p<0.001 uncorrected). Word contrast: English & French words > faces, bodies, houses, tools; face contrast: faces > bodies, houses, tools, English & French words). The majority of participants (18/21) had anterior VOTC clusters (from TAL Y > -50 to the brain stem), except the first 3 participants (SE01, SB01, SB02) affected by signal dropout above the ear canals.

**Figure S3.**
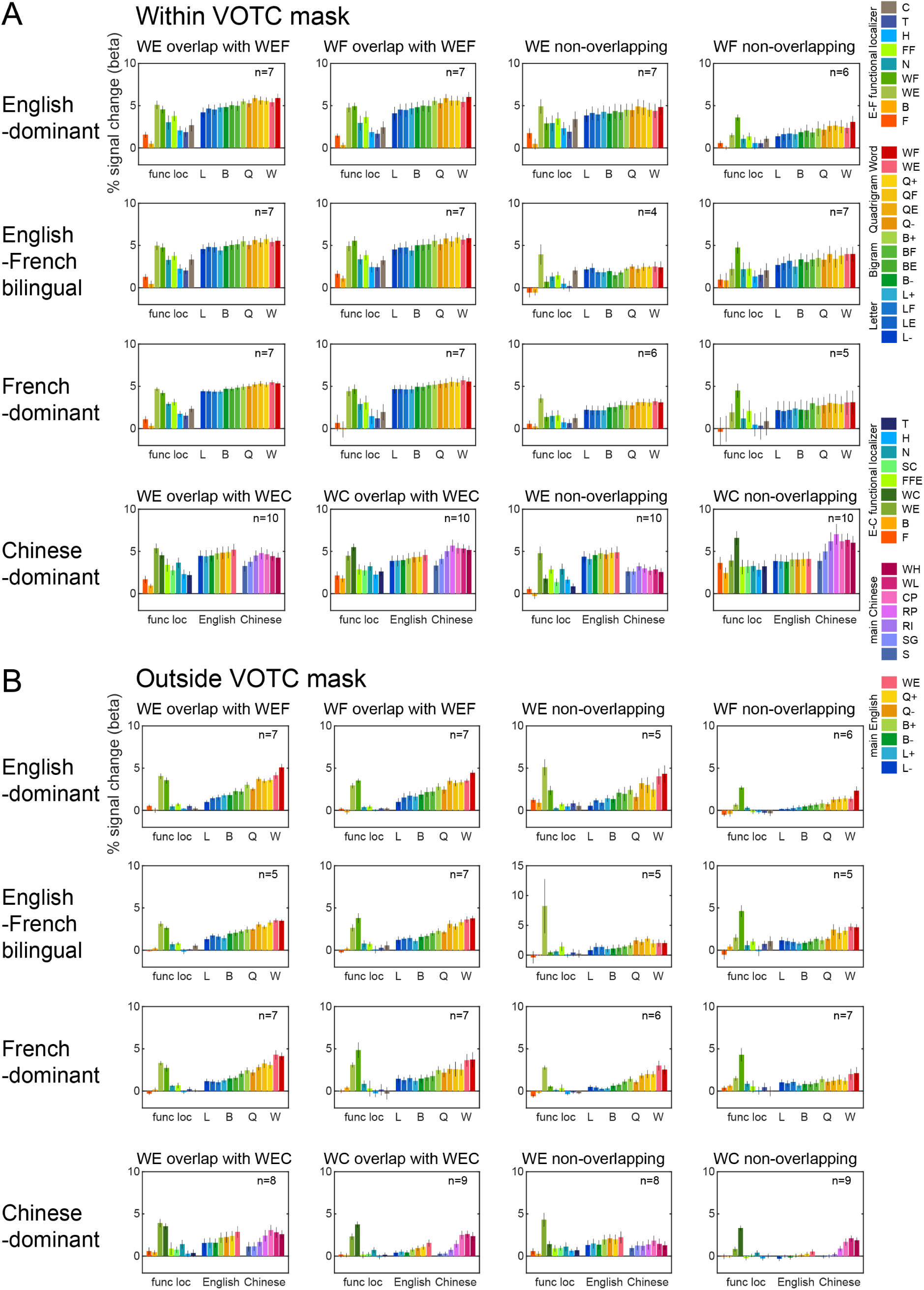
Average activity profiles for single-language-specific voxels which do or do not overlap with bilingual word-specific voxels. **A.** Voxels within the VOTC mask. **B.** Voxels outside the VOTC mask. For the English-Chinese participant (Chinese-dominant), single-language-specific voxels not overlapping with bilingual word-specific ones showed language-specificity in the activity profiles, which was not the case for any sub-groups of English-French participants. The single-language-specific conditions were abbreviated as WE, WF and WC. Error bars denote SEM across participants. The number of participants contributing to the average profiles are noted per plot.

**Figure S4.**
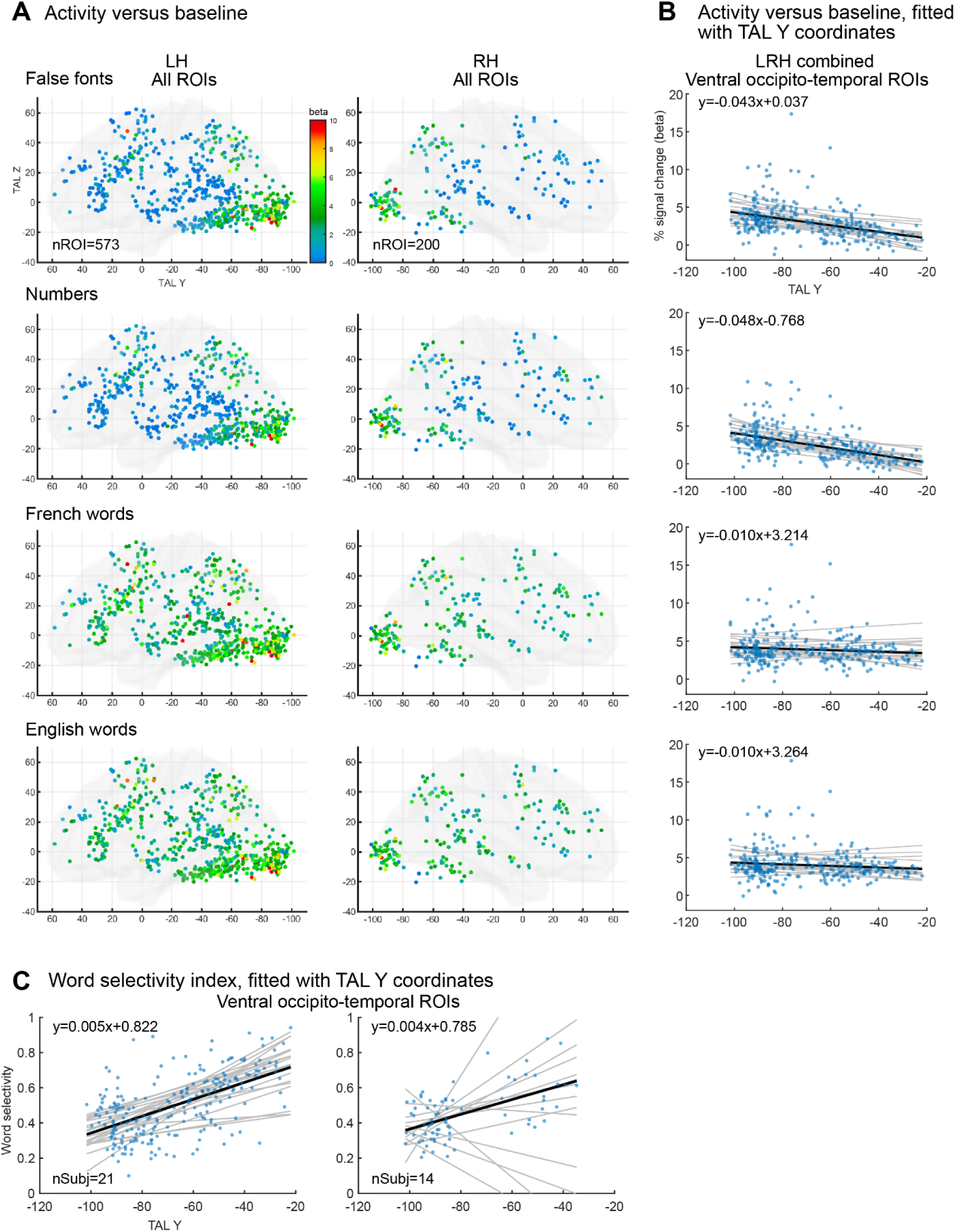
Gradient of word selectivity in word-specific ROIs across the cortex. **A.** Activity respectively evoked by false fonts, numbers, French and English words relative to baseline in the localizer. All 4 conditions yielded similar activity in occipital and posterior ventral occipito-temporal ROIs, but there was a strong selectivity for words in anterior OTS. **B**. Response amplitude in VOTC plotted against the TAL Y coordinates, corresponding to the conditions in **A** (ROIs combined across left and right hemispheres). **C**. Word selectivity index in left and right ventral ROIs, plotted against TAL Y coordinates. The word selectivity index was computed as [word activity - other activity]/[word activity + other activity] (word activity: averaged activity of English and French words; other activity: averaged activity of faces, bodies, houses, tools). If a condition with the smallest activity amplitude was negative, all conditions were padded with this absolute value, so that the activity for all conditions was zero or positive, and the resulting word selectivity index would range from -1 to 1. In both **B** and **C**, the black lines are the linear fits to all ROIs across participants (parameters in upper left corner) and the grey lines were the linear fits for individual participants.

**Figure S5.**
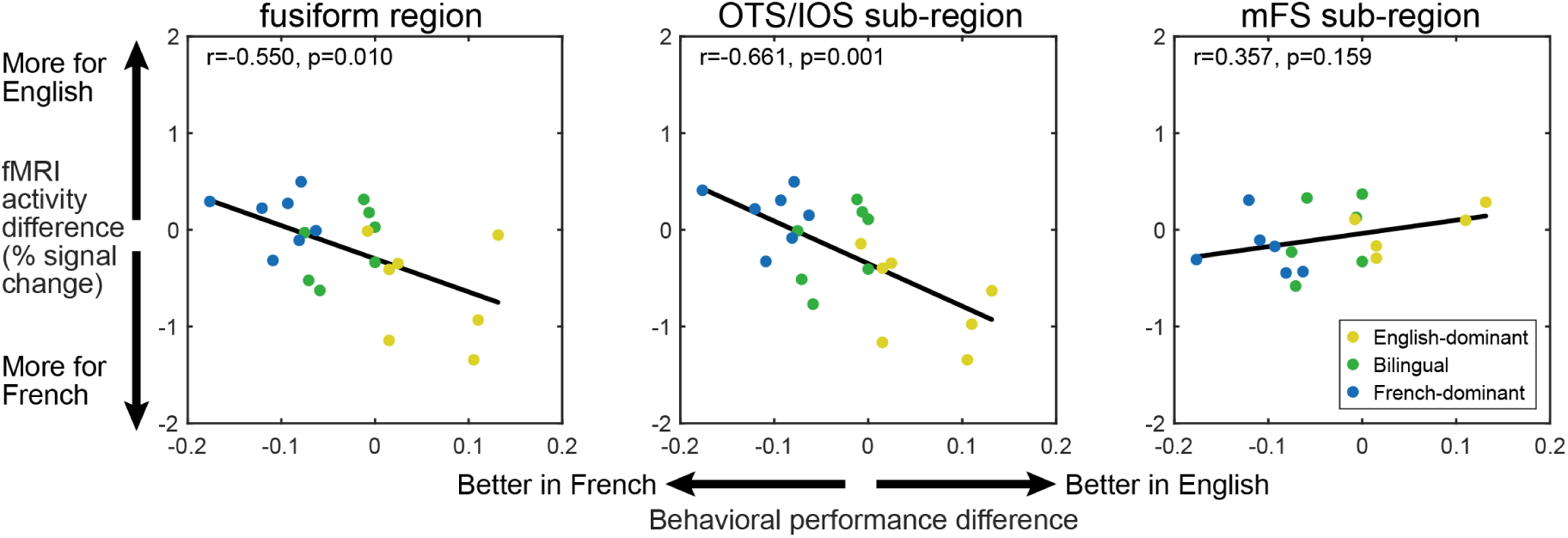
fMRI correlates of behavioral language dominance in the English-French bilingual readers. The difference in fMRI activity evoked in the fusiform region by English and by French words in the main fMRI runs (y axis) was negatively correlated with the difference in performance for those languages (x axis), indicating that the language that was less well mastered yielded a larger activity. The behavioural performance score was computed from the word counts in the one-minute reading task as [nWE - nWF] / [nWE + nWF], where nWE and nWE are the number of words read in English and French respectively.

**Figure S6.**
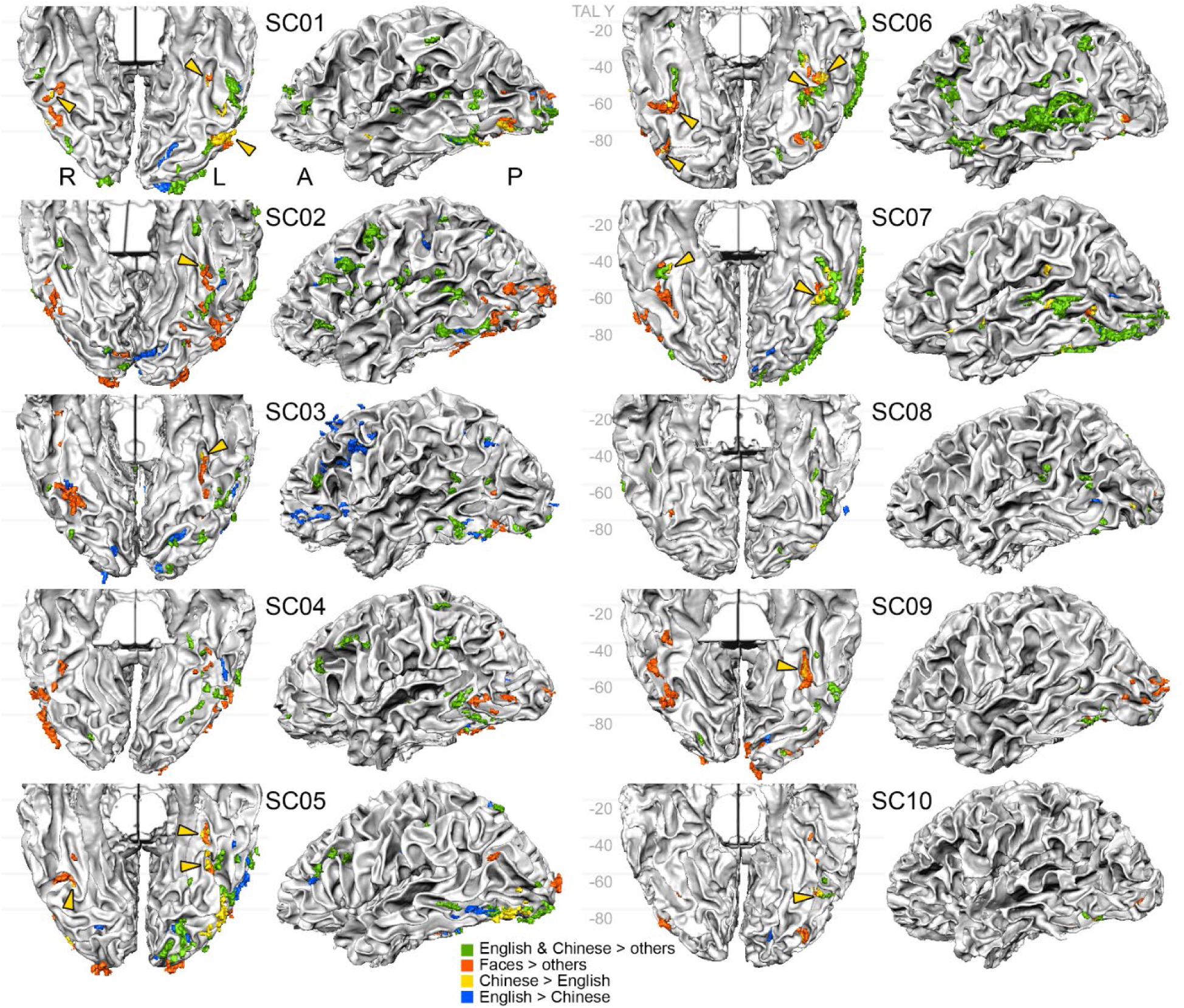
Word, face, and language-specific clusters in VOTC in each individual English-Chinese participant. (p<0.001 uncorrected). The inferior and left lateral views of individual participants show four contrast maps. Two were from the localizer: English and Chinese words > faces, bodies, houses, tools; and faces > bodies, houses, tools, English and Chinese words. Two were from the main fMRI runs: Chinese words > English words and vice-versa. For better visibility, in the inferior view, Chinese-specific clusters that are localized close to face-specific clusters are marked by yellow arrows. 8/10 participants had such clusters (participants SC04 and SC08 lack them). For the English > Chinese contrast, participant SC03 was an outlier due to too many (more than 200) activated clusters at p<0.001, and the contrast was instead plotted at p<0.0001.

**Figure S7.**
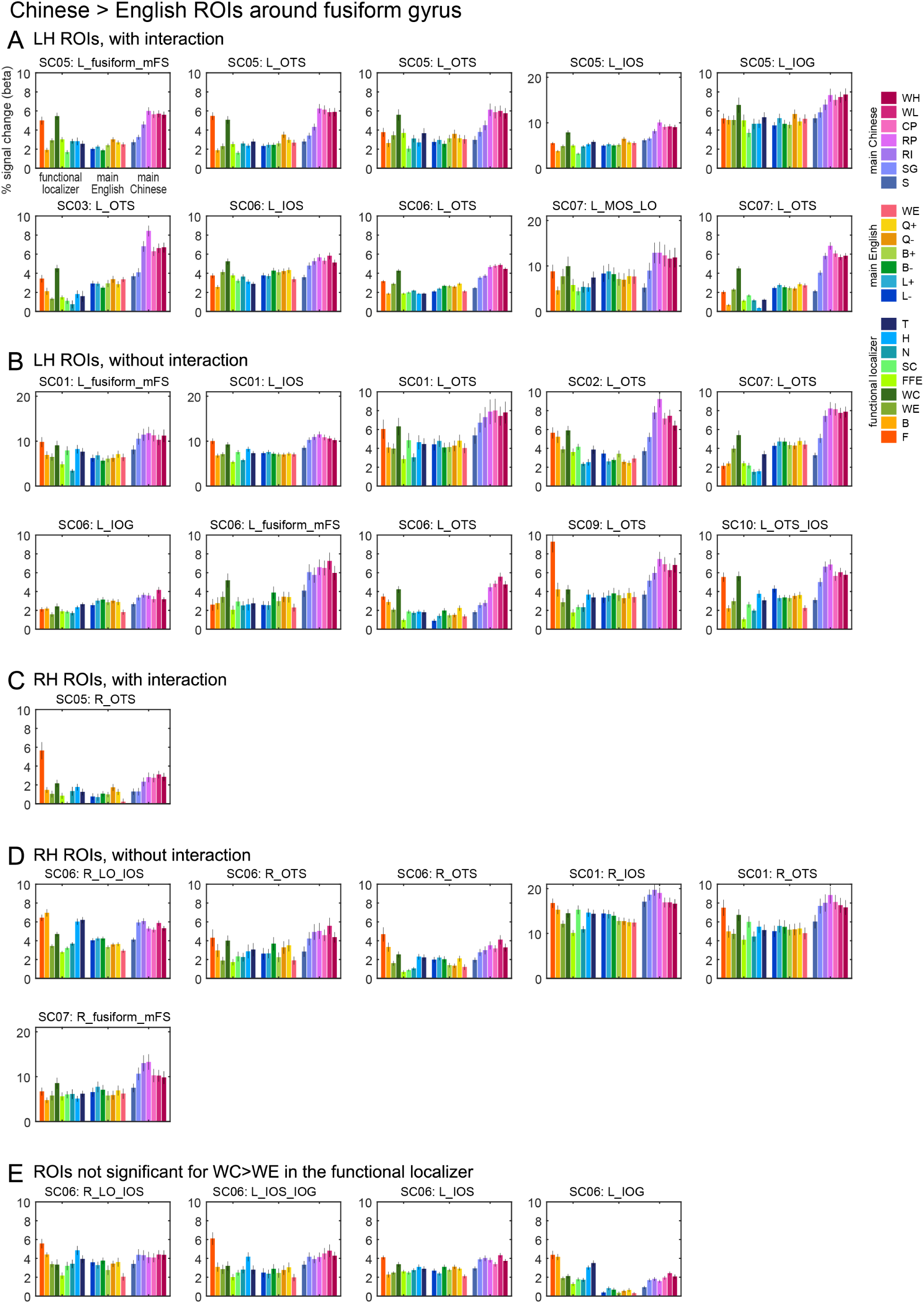
Activity profiles of all Chinese > English ROIs around the fusiform gyrus. ROIs are sorted by hemispheres, and according to: first, whether the language-specific effect in the main fMRI runs was also significant in the localizer (A-D) or not (E); second, whether the interaction (WC – WE) > (SC – FFE) was significant, indicating that the Chinese > English difference was not present for retinotopically matched control stimuli, thus not purely driven by low-level visual features associated to each language. The participant ID and ROI names are indicated on each plot.

**Figure S8.**
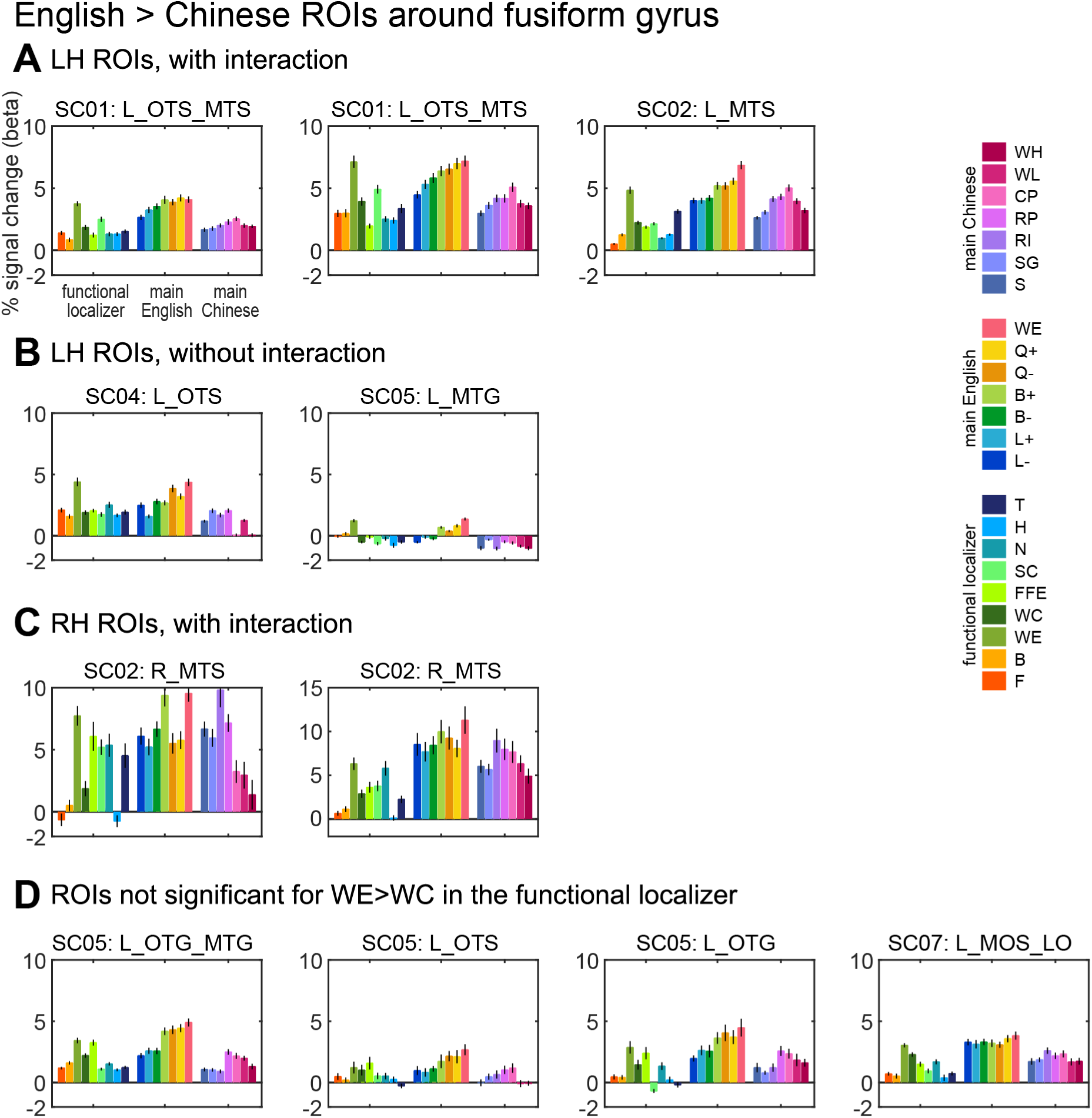
Activity profiles of all English > Chinese ROIs around the fusiform gyrus. ROIs are sorted in the same way as in figure S7.

